# Assessment of AlphaFold2 residue conformations for human proteins

**DOI:** 10.1101/2022.01.28.478137

**Authors:** Kristoffer T. Bæk, Kasper P. Kepp

## Abstract

**Motivation:** As only 35% of human proteins feature (often partial) PDB structures, the protein structure prediction tool AlphaFold2 (AF2) could have massive impact on human biology and medicine fields, making independent benchmarks of interest. We studied AF2’s ability to describe the backbone solvent exposure as an easily interpretable “natural coordinate” of protein conformation, using human proteins as test case.

**Results:** After screening for appropriate comparative sets, we matched 1818 human proteins predicted by AF2 against 7585 unique experimental PDBs, and after curation for sequence overlap, we assessed 1264 comparative pairs comprising 115 unique AF2-structures and 652 unique experimental structures. AF2 performed markedly worse for multimers, whereas ligands, cofactors and experimental resolution were interestingly not very important for performance. AF2 performed excellently for monomer proteins. Challenges relating to specific groups of residues and multimers were analyzed. We identify larger errors for lower-confidence scores (pLDDT) and exposed residues, and polar residues (Asp, Glu, Asn e.g.) being less accurately described than hydrophobic residues. Proline conformations were the hardest to predict, probably due to common location in dynamic solvent-accessible parts. In summary, using solvent exposure as a natural metric of local conformation, we quantify the performance of AF2 for human proteins and provide estimates of the expected error as a function of ligand presence, multimer/monomer status, resolution, local residue solvent exposure, pLDDT, and amino acid type. Overall performance was found to be excellent.

**Availability and Implementation:** Scripts used to perform benchmarking are available at https://github.com/ktbaek/AlphaFold.

## Introduction

The deep-learning protein-structure prediction tool AlphaFold2 (AF2)(Jumper *et al*., 2021) has generated enormous excitement in broad areas of biology and medicine (Thornton *et al*., 2021) due to its accurate prediction of protein structure from sequence, an outstanding challenge in biology (Marks *et al*., 2012). With very many protein structures still experimentally undetermined, such high-quality predicted structures are relevant to drug design, understanding protein evolution and pathogenic mutations, protein-or-peptide-based therapeutics, and structure-guided rational protein design (Jones and Thornton, 2022; Perrakis and Sixma, 2021; Skolnick *et al*., 2021; Baek *et al*., 2021; Tong *et al*., 2021; Monzon *et al*., 2022; Tsaban *et al*., 2022).

AF2 combines multi-sequence analysis with advanced structural pattern recognition that accounts for intra-protein epistasis (amino acid correlations in space not captured by the sequences directly that modify each amino acid’s structure and substitution likelihood (Mumenthaler and Braun, 1995; Ortiz *et al*., 1998)). With additional vast amounts of structural and sequence data available for training, such deep learning of the 3D residue correlations has created unprecedented descriptive performance (Xu, 2019; Skolnick *et al*., 2021; Perrakis and Sixma, 2021). As independent assessments emerge, the high quality of AF2 seems confirmed (Akdel *et al*., 2021). Performance is naturally best for structured parts, but this is not critical since dynamic coils and loops in solution are not likely to be well modeled by any static structure, and would be sensitive to changes in environment. The method’s training data and thus performance is necessarily better for types of protein structures that are best covered empirically, and work remains to determine the range of confidence not on a per residue level (as AF2 estimates directly) but also for structurally rare proteins with less training set coverage, membrane proteins, interface structures of multimeric proteins, proteins with cofactors or modifications, and intrinsically disordered proteins (Perrakis and Sixma, 2021).

Among these challenges, the local solvent exposure of a residue, both in monomers and in monomer interfaces of multimeric proteins, is of particular importance (Evans *et al*., 2021). The solvent exposure provides important information on protein compactness, local residue conformation, and chemically relevant features of the site, e.g. in relation to prediction of mutation rates in protein evolution and protein stability effects, which directly depend on solvent exposure (Bloom *et al*., 2006; Zhou *et al*., 2008; Caldararu *et al*., 2021). Being able to describe residue solvent exposure via AF2 would thus mean potentially better estimates of structure-informed evolution rates (substitution frequencies) and protein stability effects (Caldararu *et al*., 2021) making this parameter a natural (and chemically interpretable) coordinate in contrast to e.g. XYZ coordinates of the residues and associated root-mean-square-deviations.

Critical assessment of AF2 requires curated and unbiased test sets that separate multimer and monomer proteins, account for the data-coverage bias (bias in training and test sets towards certain types of proteins) and quantify performance on a per residue-level for a focused and application. Below we attempt such an analysis for human proteins as described by AF2 recently (Tunyasuvunakool *et al*., 2021).

## Methods

### Structures used in the test set

The AlphaFold predictions for 23,391 proteins of the *Homo sapiens* reference proteome (UniProt proteome ID UP000005640) were downloaded from https://AlphaFold.ebi.ac.uk/download on September 16, 2021 (version v1)(Varadi *et al*., 2022). In addition, PDB files for all human protein structures that have been solved by X-ray crystallography with a resolution ≤ 2.0 Å (giving a total of 19,710 PDB structures) were retrieved from the protein data bank(wwPDB consortium, 2019; Berman *et al*., 2000) (https://www.rcsb.org) in .pdb format on October 1, 2021. These structures were then further curated into relevant non-biased test sets and subsets to account for performance confounders, as described below.

### Matching AF2 structures to experimental structures

To compare protein structures predicted by AF2 to protein structures solved experimentally, we first needed to match each experimental structure to the AF2 prediction of the protein. This was done in a two-step process in which each AF2 structure was first paired to one or more experimental PDBs by matching UniProt numbers, if possible, and then the correct experimental chain structure was identified by sequence alignment as outlined in **Figure 1**.

**Figure 1:**
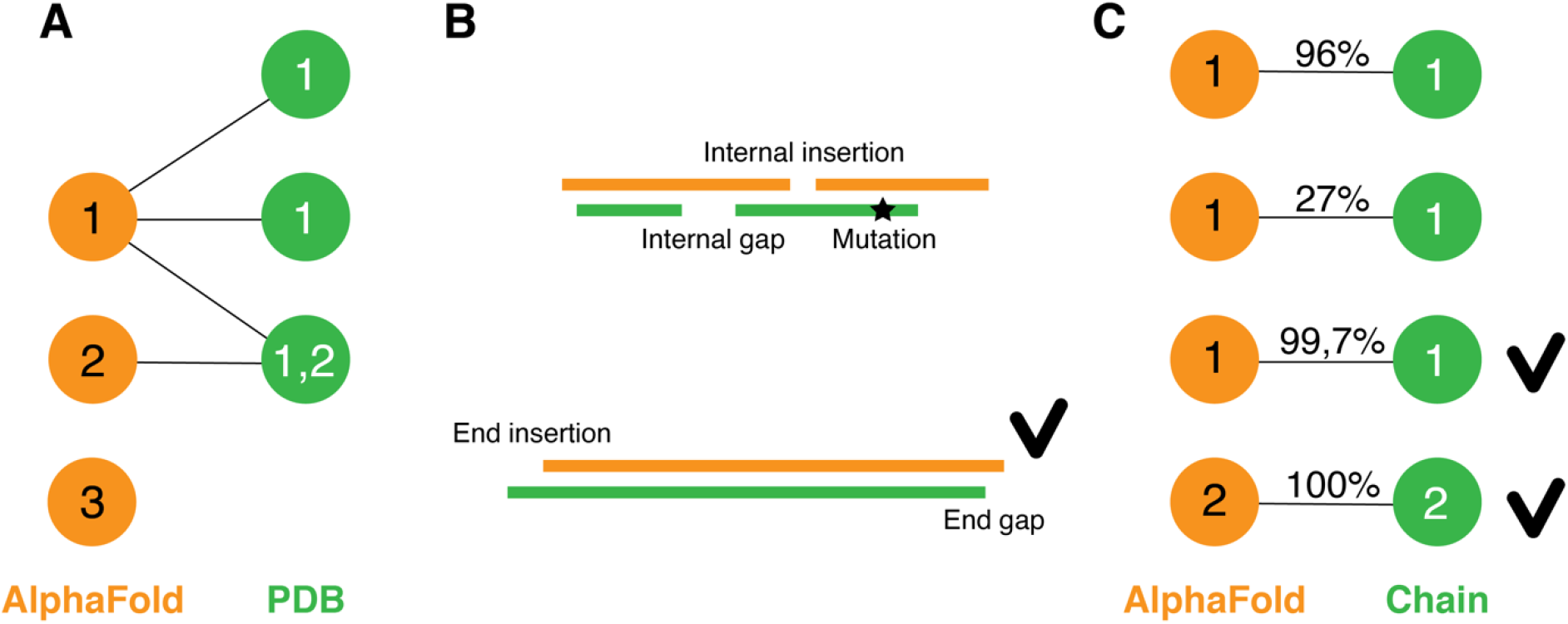
Protocol for matching AlphaFold structures to experimental structures. **(A)** Each AlphaFold structure was matched to experimental PDBs by UniProt number. AlphaFold structures can match none, one or more PDBs. PDBs can match one or more (i.e., heteromultimer) AlphaFold structures. **(B)** Examples of alignments of AlphaFold and experimentally solved sequences (from the ATOM field of the PDB file). Pair alignments with internal gaps, insertions or mutations (upper panel) were discarded, while gaps or insertions at ends were tolerated. **(C)** Four different pairs with different degrees of sequence overlap (exemplified by indicated percentages). Numbers inside circles indicate UniProt ID. Checkmarks indicate pairs that are part of the final dataset.

To match by UniProt number, the unique UniProt number of each AlphaFold structure was used to search through the retrieved PDBs described above. This resulted in 3930 AF2-structures being matched to 18,389 unique experimental PDBs. Some AF2-structures were matched to several PDBs, and some PDBs were matched to more than one AF2-structure, if the PDB represented the structure of a heteromultimer (**Figure 1A**).

To 1) identify the correct experimental chain structures for each AF2-structure and 2) assess how much of the protein chain was actually solved in the crystal structure, the resolved amino acid sequences were extracted from the PDB files using the residue information in the ATOM field, not the SEQRES field, of the PDB. The resolved amino acid sequence of each chain was then aligned by pairwise alignment to its UniProt-paired amino acid sequence. Experimental sequences that had either internal gaps, internal insertions, or mutations compared to the UniProt (and thus AF2) sequence were excluded from further analyses (**Figure 1B**). This resulted in 16,036 pairs, each pair consisting of one AF2 structure and one experimental chain structure (in the following referred to as data pairs).

The set of comparable pairs resulting from this protocol comprised 2249 unique AF2 structures, 12,520 unique experimental chains, and 7585 unique experimental PDBs. In the AF2 dataset, proteins longer than 2700 amino acid residues are split into 1400 amino acid long, overlapping structure fragments that may all match the same experimental chain given the described matching protocol (Varadi *et al*., 2022). Therefore, in this dataset, the number of data pairs exceed the number of experimental chains, and the number of unique AF2 structures exceed the number of unique human proteins (1818). This dataset thus emphasizes the experimental structures with either 100% sequence identity to the AF2 sequence or with gaps or insertions only in the beginning or end of the sequence compared to the AF2 sequence (**Figure 1C**). We consider this curation important, since experimental PDB structures with missing or modified parts could erroneously affect performance.

Next, we reduced this dataset further to include only comparable pairs of structures for which the overlap between sequences comprised >99% of the experimental and the AF2/canonical sequence length. The proportion of sequence overlap was calculated as the length of overlap divided by the length of sequence, and the data pairs were restricted to those whose length of overlap constituted >99% of both the UniProt sequence and the experimental sequence. This last exclusion step resulted in a final dataset consisting of 1264 data pairs comprising 115 unique AF2 structures and 652 unique PDBs (**Table S1**). The length of overlapping sequence in the data pairs ranged from 68 to 888 residues with an average length of 197 residues. The above procedure was performed with Python using BioPython v.1.78 modules (Cock *et al*., 2009).

### Calculation of solvent accessible surface

The relative solvent accessible area (RSA) is a descriptor of local conformation that is more easily interpretable than XYZ coordinates. RSA was computed using the open-source software FreeSASA available as a Python module using the classifier configuration option “naccess” (Mitternacht, 2016). We only used the solvent accessibility of the backbone (including the C_α_ atom) without the side-chain, as the prediction of side-chain orientation will have less certainty (Jumper *et al*., 2021) and confound the analysis unfairly, as side chain conformations are inherently dynamic and not necessarily important to be accurately described vs. a crystal structure (the solution dynamics of surface residue side chains can be studied by other non-static methods such as molecular dynamics and NMR).

The RSA is defined as the ratio of the absolute solvent accessible area of a residue divided by that within an Ala-X-Ala tripeptide, where X is the residue studied. For each data pair, the RSA values for each residue in the AF2-structure (RSA_AF_) and the experimental structure (RSA_Exp_), respectively, were computed with FreeSASA. Then, each residue in the overlapping portion of the two sequences were matched using residue number and residue type, and the RSA_AF_ and RSA_Exp_ values were assigned to each common residue. Because the numbering of residues in the experimental PDBs may not be consistent with the numbering in the AF2-structures, a temporary common number was assigned to each residue to allow unambiguous matching of residues. The first and last residue in each structure, both AF2 and experimental, were excluded from the analyses due to the risk of overestimated RSA values.

### AlphaFold per-residue confidence values

AlphaFold predicts residue coordinates with different confidence using a per-residue estimate called predicted lDDT-C_α_ (pLDDT), which has numbers on a scale from 0 to 100 (Mariani *et al*., 2013; Jumper *et al*., 2021). pLDDT values for each residue are stored in the B-factor fields of PDB files (Varadi *et al*., 2022), and to assess the effect of pLDDT, the B-factor fields were extracted from the PDB files using BioPython v.1.78 modules.

### Ligands

Experimental structures can have non-covalently bound ligands such as metabolites, inhibitors, drugs, cofactors, and ions (Sen *et al*., 2014). To assess the effect of such co-solutes on performance, the experimental structures in the dataset were classified according to ligands. Ligand information was extracted from the HET field in the PDB files using Python, and experimental structures with no ligands or only water or the small ions Na^+^, Cl^-^, K^+^, F^-^, Li^+^, Br^-^, I^-^, SO_4_^2-^, or PO_4_^3-^ (HET codes HOH, NA, CL, K, F, LI, BR, I, SO4, and PO4) were classified as having no ligands.

### Dataset subsets

The dataset (**Table S1**) was divided into six disjointed subsets based on 1) proportion of sequence overlap (100 %, or >99% and <100%), 2) resolution of the experimental structure (<= 1.5 Å, or >1.5 Å and <=2.0 Å), and 3) whether the experimental structure is a monomer. Since the number of pairs with 100% sequence identity was small, this group was not split based on resolution. The six data subsets have the following characteristics: 1) 100% sequence identity, ≤ 2.0 Å, and monomer; 2) 100% sequence identity, ≤ 2.0 Å, and multimer; 3) > 99% and <100% sequence identity, ≤ 1.5 Å resolution, and monomer; 4) > 99% and <100% sequence identity, ≤ 1.5 Å resolution, multimer; 5) > 99% and <100% sequence identity, > 1.5 Å and ≤ 2.0 Å resolution, and monomer; 6) > 99% and <100% sequence identity, > 1.5 Å and ≤ 2.0 Å resolution, and multimer.

### Identification of chain-interface residues in multimers

In multimeric proteins, residues from one chain may be in close contact to residues from another chain, potentially affecting these residues’ solvent accessibility. To analyze the effect of these chain-interface residues, all experimental multimeric structures in the dataset were analyzed with BioPython to identify chain-interface residues in the chains that were paired to an AF2-structure. For each atom of each residue in the chain, the distance to atoms belonging to other chains in the structure was calculated, and residues that contained an atom that were less than 3.5 Å from atoms in other chains were classified as chain-interface residues.

## Results and discussion

### Performance of AF2 as measured by relative solvent accessibility

There are multiple ways of assessing a structural prediction by AF2, including various root-mean-square-deviations between the experimental and predicted structures, including backbone atoms, and also the robustness of the predicted structure’s ability to predict important protein properties (Akdel *et al*., 2021). In this work we emphasize RSA as a directly interpretable endpoint of evaluation, or a “natural” coordinate” of each residue in the context of the protein fold with direct implication for function and evolution, as opposed to residual XYZ coordinate variations.

Since only 35% of human proteins feature (often partial) PDB structures (Tunyasuvunakool *et al*., 2021), AF2 could have massive impact for understanding human diseases in protein-structural context, making independent benchmarks focusing on the human proteome relevant. We compared AF2 predictions of proteins in the human proteome to their experimentally solved structures. We limited the comparison to structures solved by X-ray crystallography with a resolution of 2.0 Å or better. Of the 23,391 human protein chains predicted by AF2 (Tunyasuvunakool *et al*., 2021; Varadi *et al*., 2022), 115 were successfully matched to 1264 experimental chain structures from 652 PDBs. In order to avoid erroneous effects, we excluded experimental structures with internal gaps, insertions or mismatches (mutations) compared to the UniProt sequence (i.e. only 100% sequence identity or experimental structures with gaps or insertions only in the beginning or end of the sequence) (**Figure 1B**). For pairs of AF2- and experimental structures with less than 100% sequence identity, we only included overlaps between sequences of more than 99% of the sequence length (**Figure S1**). The final dataset included 1264 pairs comprising 115 unique AF2-structures and 652 unique PDBs (**Table S1**). The overlapping sequences ranged from 68 to 888 residues with an average length of 197 residues.

We then calculated the RSA for each residue backbone (including the C_α_ atom) in both the AF2-generated (RSA_AF_) and experimental structures (RSA_Exp_), and compared the RSA values for each matched residue in each structure pair. Because AF2 generally predicts side-chain coordinates with less accuracy than backbone coordinates (Jumper *et al*., 2021), we compared only backbone RSA values. Scatter plots showing the correlation between RSA_AF_ and RSA_Exp_ for each of the 1264 analyzed data pairs are shown in **Figure S2.** In order to understand AF2’s performance in detail, we divided the data pairs into six non-overlapping groups based on 1) length of sequence overlap, 2) resolution of experimental structure, and 3) monomer vs. multimer structures. We compared the RSA for each matched residue in the structure pairs belonging to the six groups (**Figure 2A**). For each group, we calculated the mean absolute error (MAE), the mean signed deviation (MSD), and the standard deviation of the absolute errors (SA_Abs_, **Table 1**).

**Figure 2:**
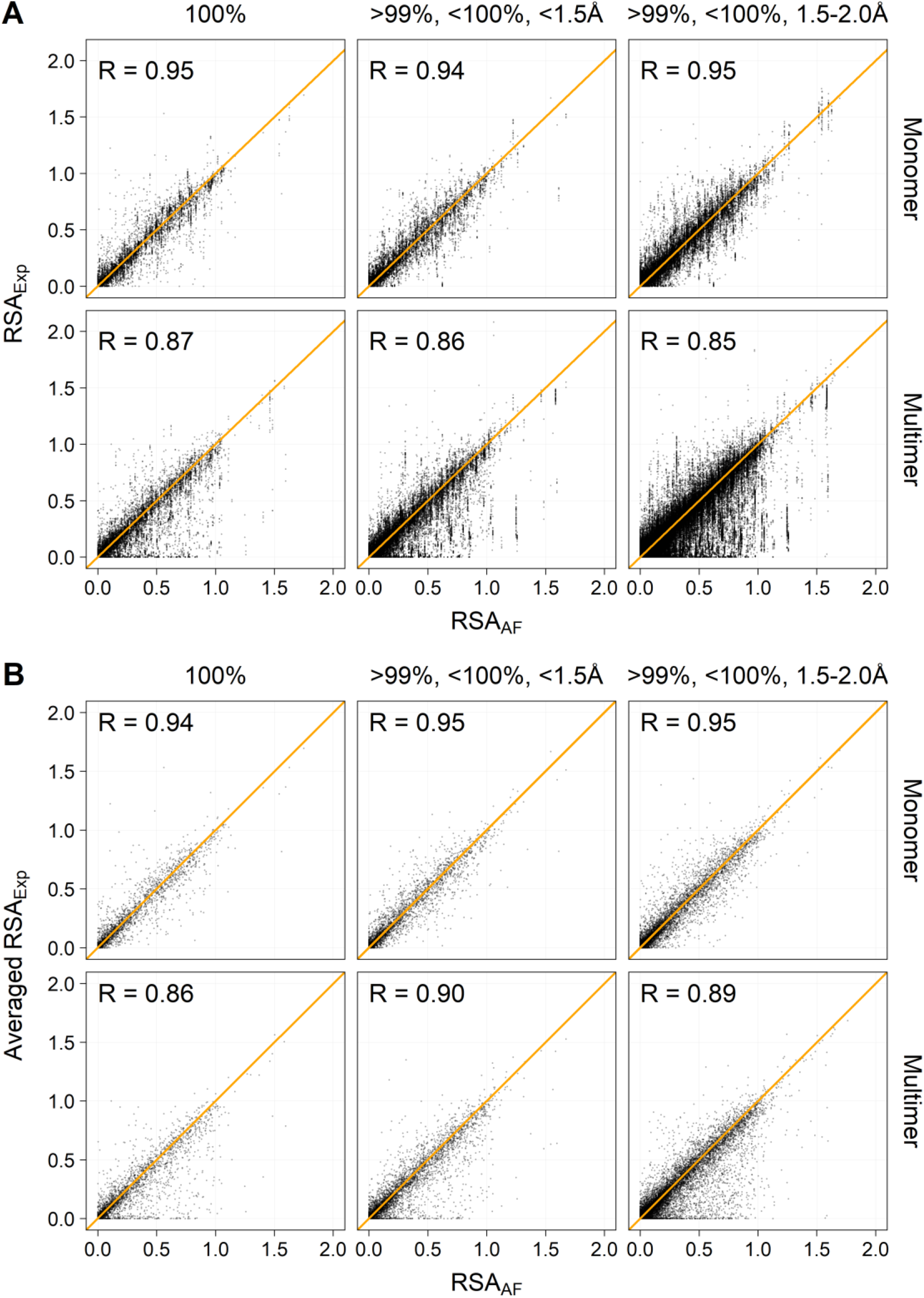
Experimental vs. AF2 RSA values. Each panel shows results grouped based on sequence overlap, resolution of the experimental structure, and monomer-multimer status of the experimental structure (indicated above and to the right). The six groups are disjointed from each other. The orange line represents the ideal where the RSA_AF_ are equal to RSA_Exp_. The correlation coefficient R is indicated for each group. **(A)** Each dot represents a residue belonging to an Data pair. **(B)** Each dot represents the average of experimental residues belonging to the same AF2-structure and the same overlap-, resolution- and monomer/multimer group.

**Table 1.**
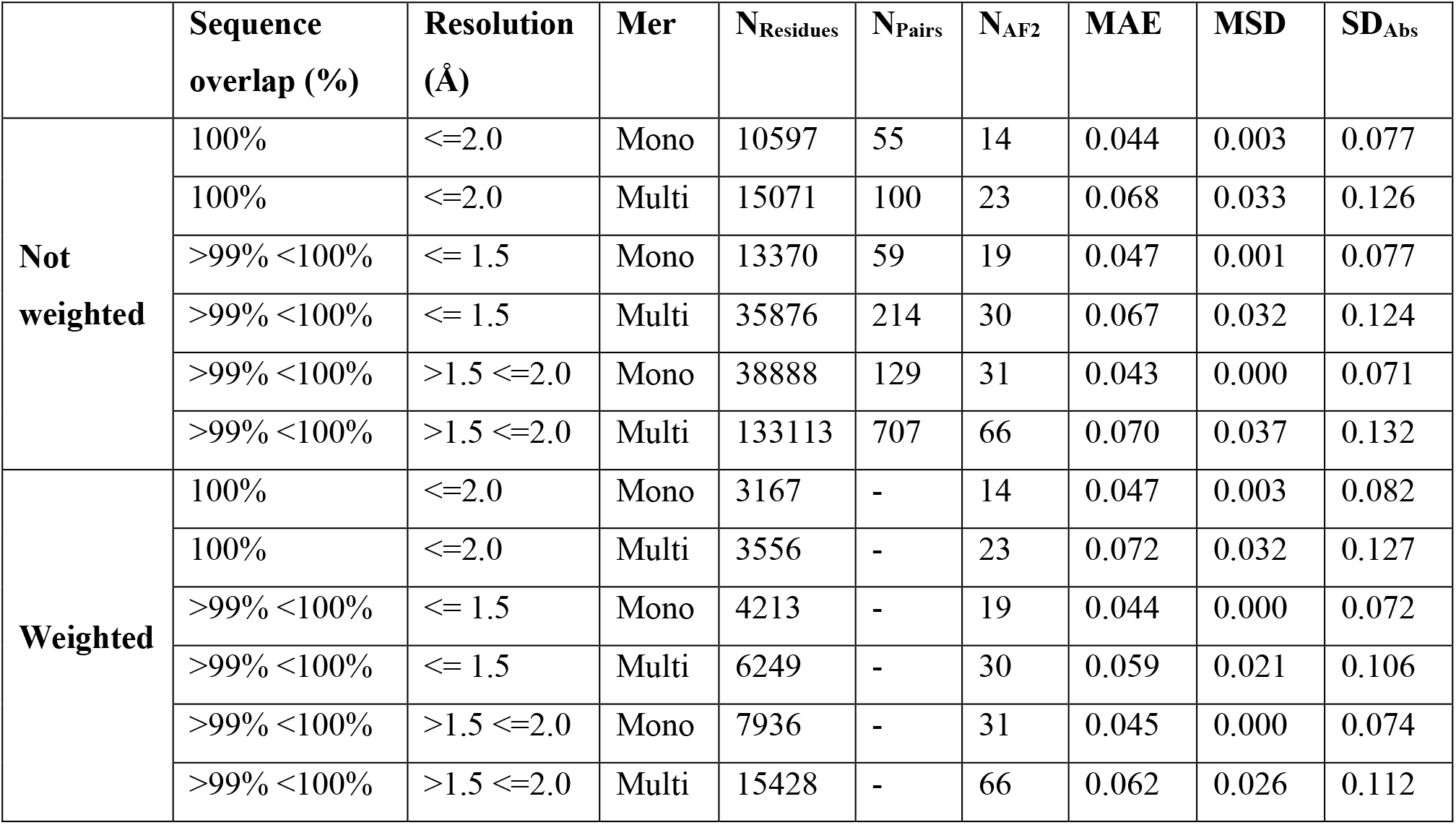
Characteristics of the six groups of data pairs.

| | Sequence overlap (%) | Resolution (Å) | Mer | $N_{Residues}$ | $N_{Pairs}$ | $N_{AF2}$ | MAE | MSD | $SD_{Abs}$ |
| --- | --- | --- | --- | --- | --- | --- | --- | --- | --- |
| **Not weighted** | 100% | $\leq 2.0$ | Mono | 10597 | 55 | 14 | 0.044 | 0.003 | 0.077 |
| | 100% | $\leq 2.0$ | Multi | 15071 | 100 | 23 | 0.068 | 0.033 | 0.126 |
| | >99% <100% | $\leq 1.5$ | Mono | 13370 | 59 | 19 | 0.047 | 0.001 | 0.077 |
| | >99% <100% | $\leq 1.5$ | Multi | 35876 | 214 | 30 | 0.067 | 0.032 | 0.124 |
| | >99% <100% | $>1.5 \leq 2.0$ | Mono | 38888 | 129 | 31 | 0.043 | 0.000 | 0.071 |
| | >99% <100% | $>1.5 \leq 2.0$ | Multi | 133113 | 707 | 66 | 0.070 | 0.037 | 0.132 |
| **Weighted** | 100% | $\leq 2.0$ | Mono | 3167 | - | 14 | 0.047 | 0.003 | 0.082 |
| | 100% | $\leq 2.0$ | Multi | 3556 | - | 23 | 0.072 | 0.032 | 0.127 |
| | >99% <100% | $\leq 1.5$ | Mono | 4213 | - | 19 | 0.044 | 0.000 | 0.072 |
| | >99% <100% | $\leq 1.5$ | Multi | 6249 | - | 30 | 0.059 | 0.021 | 0.106 |
| | >99% <100% | $>1.5 \leq 2.0$ | Mono | 7936 | - | 31 | 0.045 | 0.000 | 0.074 |
| | >99% <100% | $>1.5 \leq 2.0$ | Multi | 15428 | - | 66 | 0.062 | 0.026 | 0.112 |

The MAE, the MSD, and the spread of absolute errors are very dependent on the monomer-multimer status of the experimental structure, but not on the length of the sequence overlap or the resolution of the experimental structure (**Table 1**). The higher MAE, MSD, and spread of errors for multimeric structures is likely caused mainly by residues that have low RSA values in the experimental structure but high RSA values in the AF2 structure (**Figure 2A,** lower panels). This illustrates why caution should be exercised when deducing surface structure for multimeric proteins based on AF2 predictions for a single monomer, especially at the interfaces. After establishing this, we restricted further analysis mainly to monomer comparisons.

There was very large variation in the number of experimental structures matching individual AF2-structures (**Table S1**). Some AF2-structures matched only one experimental structure whereas a few (such as hemoglobin subunit alpha, UniProt ID P69905) matched hundreds of experimental structures (**Figure S3**). This variation may potentially skew the error metrics (MAE, MSD and SD_Abs_) towards those of highly represented structures. To adjust for this bias, we calculated the average RSA_Exp_ for each residue that were paired to the same AF2 structure if the pair belonged to the same overlap-, resolution- and monomer/multimer group described above **(Figure 2B)**. Similar to the first analysis (**Table 1**, upper half), the absolute and signed deviations calculated from the averaged RSA_Exp_ values resulted in MAE, MSD, and SD_Abs_ values that depended highly on the monomer-multimer status, but not on the length of sequence overlap or resolution of the experimental structure (**Table 1**, lower half).

### The effect of non-covalently bound ligands on prediction performance

Many experimental protein structures in the PDB have non-covalently bound ligands such as metabolites, inhibitors, drugs, cofactors, ions, and solvent (Sen *et al*., 2014). We therefore tested whether the observed differences between RSA_AF_ and RSA_Exp_ correlated with the presence of ligands, by first classifying the experimental structures in the dataset as being with or without ligands, and then measuring the RSA prediction accuracy separately on these two subsets. For this analysis, water or the small ions, Na^+^, Cl^-^, K^+^, F^-^, Li^+^, Br^-^, I^-^, SO_4_^2-^, or PO_4_^3-^ were not counted as ligands. We found no apparent correlation between the MAE, the MSD, or SD_Abs_ and the presence of ligands (excluding water and small ions, **Table 2**); the performance for the monomer structures was very similar regardless of ligand presence. We therefore continue our analysis below without separating out ligand-containing structures.

**Table 2.** The effect of ligands in the experimental structure. Ligands are heteromolecules that are not H_2_O, Na^+^, Cl^-^, K^+^, F^-^, Li^+^, Br^-^, I^-^, SO_4_^2-^, or PO_4_^3-^.

| Sequence overlap (%) | Resolution (Å) | Mer | Ligands | $N_{Residues}$ | $N_{Pairs}$ | $N_{AF2}$ | MAE | MSD | $SD_{Abs}$ |
| --- | --- | --- | --- | --- | --- | --- | --- | --- | --- |
| 100 | $\leq 2.0$ | Mono | No | 1306 | 7 | 6 | 0.044 | 0.000 | 0.079 |
| 100 | $\leq 2.0$ | Mono | Yes | 9291 | 48 | 10 | 0.044 | 0.003 | 0.077 |
| 100 | $\leq 2.0$ | Multi | No | 1624 | 11 | 4 | 0.080 | 0.043 | 0.153 |
| 100 | $\leq 2.0$ | Multi | Yes | 13447 | 89 | 21 | 0.066 | 0.032 | 0.122 |
| >99 <100 | $\leq 1.5$ | Mono | No | 558 | 4 | 3 | 0.040 | 0.005 | 0.065 |
| >99 <100 | $\leq 1.5$ | Mono | Yes | 12812 | 55 | 19 | 0.047 | 0.001 | 0.077 |
| >99 <100 | $\leq 1.5$ | Multi | No | 603 | 4 | 4 | 0.075 | 0.022 | 0.139 |
| >99 <100 | $\leq 1.5$ | Multi | Yes | 35273 | 210 | 26 | 0.067 | 0.032 | 0.124 |
| >99 <100 | $>1.5 \leq 2.0$ | Mono | No | 3980 | 25 | 7 | 0.044 | 0.002 | 0.075 |
| >99 <100 | $>1.5 \leq 2.0$ | Mono | Yes | 34908 | 104 | 26 | 0.043 | -0.001 | 0.071 |
| >99 <100 | $>1.5 \leq 2.0$ | Multi | No | 8165 | 39 | 12 | 0.070 | 0.039 | 0.137 |
| >99 <100 | $>1.5 \leq 2.0$ | Multi | Yes | 124948 | 668 | 60 | 0.070 | 0.037 | 0.132 |

### Larger errors for lower-confidence scores and exposed residues

In order to determine the factors that affect the accuracy of RSA predicted by AF2, we first hypothesized that residues with high prediction confidence or residues buried in the core of the protein were predicted more accurately. To test this hypothesis, we only analyzed data pairs in which the experimental structure is a monomer. AF2 produces an estimate of confidence for each residue, predicted lDDT-C_α_ (pLDDT) based on the Local Distance Difference test, on a scale from 0-100, where 100 indicates the highest possible confidence (Mariani *et al*., 2013; Jumper *et al*., 2021; Tunyasuvunakool *et al*., 2021). Regions with pLDDT > 90 are expected to be modelled to high accuracy, whereas regions with pLDDT < 50 are most likely either unstructured or only structured as part of a complex (Varadi *et al*., 2022). We hypothesized that the confidence with which AF2 predicts a residue’s coordinates correlates with the ability to correctly predict the RSA of a residue. If so, the MAE or the SD_Abs_ between RSA_AF_ and RSA_Exp_ would be lower for residues with high pLDDT values and *vice versa*.

To test this hypothesis, we divided the residues of all AF2-structures that were matched to a monomeric experimental structure into pLDDT bins as shown in **Figure 3A**, and calculated the MAE, MSD, SD_Abs_, and SD_Signed_ for each group (**Table S2**). As can be seen in **Figure 3A** (and **Figure S4** and **S5**), the MAE and the SD_Abs_ are indeed dependent on pLDDT with both values increasing substantially for residues with <90 pLDDT scores. The MSD is less dependent on pLDDT, meaning that there is no tendency for RSA_AF_ to be either under- or overestimated for residues with low pLDDT values.

**Figure 3.**
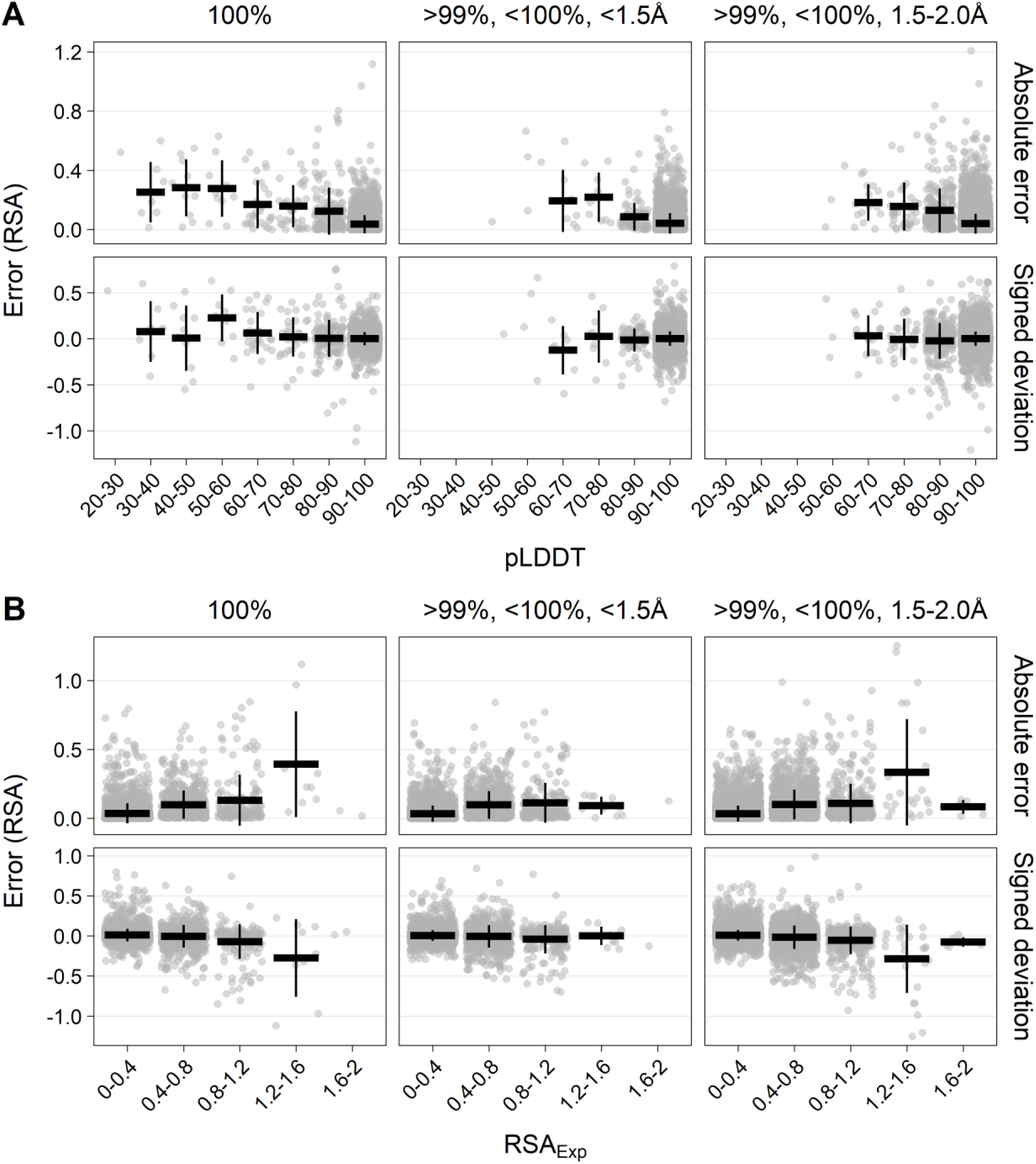
Error vs pLDDT and experimental RSA. Data pairs are grouped by length of sequence overlap and resolution of the experimental structure. Only monomeric structures included. Gray circles indicate the error (absolute error or signed deviation) calculated from the average RSA_Exp_ for each unique AF2-structure belonging to the same overlap-, resolution- and monomer/multimer group. The mean errors for each pLDDT or RSA_Exp_ group are shown as horizontal black bars and one standard deviation (SD) of the mean errors shown as vertical black bars. **(A)** Error as a function of pLDDT. The pLDDT values are grouped in 10 percent-point bins. **(B)** Error as a function of RSA_Exp_. The experimental RSA values are grouped in 0.4 RSA bins. Data for this plot are shown in **Table S2** and **Table S3**.

Next, to test if the ability of AF2 to predict RSA depended on experimentally determined solvent accessibility of the residues (RSA_Exp_), we divided the residues of all AF2-structures that were matched to a monomeric experimental structure into RSA bins, as shown in **Figure 3B**, and calculated the MAE, MSD, SD_Abs_, and SD_Signed_ for each group (**Table S3**). As can be seen in **Figure 3B** (and **Figure S6** and **S7**), the MAE and the SD_Abs_ are dependent on the experimental RSA values: Residues with low RSA, i.e. buried residues were predicted more accurately by AF2 than surface exposed residues. There is a tendency for the MSD to be more negative for high RSA residues, meaning that AF2 tends to underestimate the RSA for more surface exposed residues. The observation that errors are larger for exposed residues, and for residues with lower confidence, is of course correlated and expected, but the numbers in **Figure 3** gives an estimate of the magnitude of errors that can be expected for each class of residue. The outliers are relatively few in both cases, and the tendency of many outliers at low exposure or high confidence relates to the fact that there are many data points in total of these types.

### Larger errors for polar residues and proline

The accuracy of AF2-predicted RSA could also depend on amino acid type. In order to understand this, the same analysis as above (**Figure 3**) was performed but with all groups combined, and errors divided into amino acid type (**Figure 4**). The effect of amino acid type was remarkably large, more than 100% even after averaging errors across all pair comparisons. The best-described amino acids tend to be isoleucine, leucine, methionine, phenylalanine, tryptophan, cysteine, and valine, which are hydrophobic. The worst-described amino acids are polar, such as aspartate and glutamate, lysine, asparagine, and serine. However, proline is clearly hardest to predict. These observations can be explained by polar residues and proline being more often located in less-well-described surface areas of the proteins, i.e., with a correlation to the pLDDT/RSA errors in **Figure 3**.

**Figure 4.**
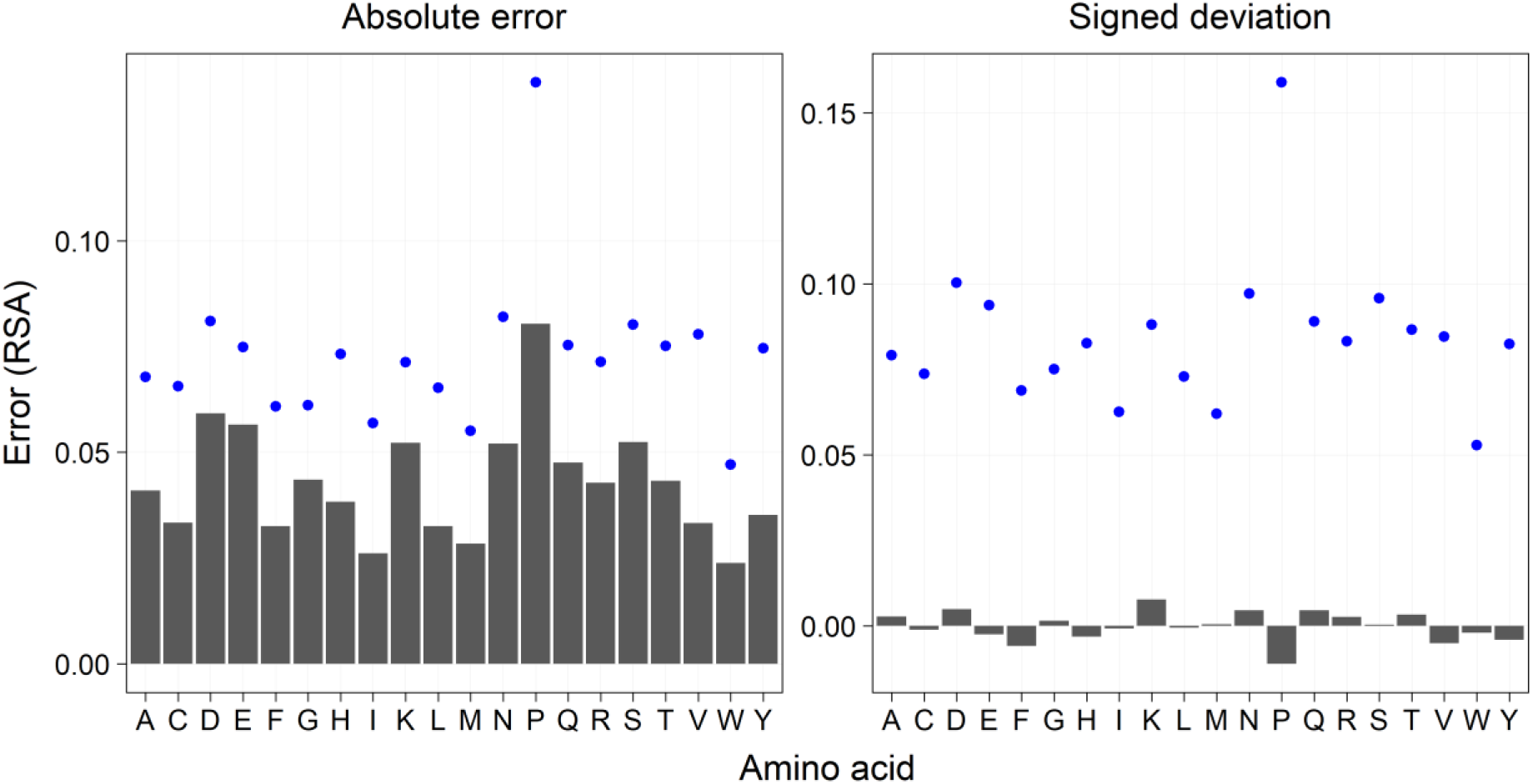
Error vs amino acid type. Left: MAE. Right: MSD (RSA_AF_ – RSA_Exp_). Only pairs with monomeric experimental structures included. The standard deviations are shown as blue dots.

While expected, **Figure 3** and **Figure 4** quantify the expected errors in a prediction for each type of residue, which is of interest to applications. Despite these errors, it is encouraging to see that the overall magnitude of the MAEs is in the range of 0.02-0.08. With RSA-values averaging to ~0.2, i.e., the expected error in RSA from an AF2 prediction is typically of the order of 20%, which we consider a relevant “natural” measure of the conformational uncertainty. A corresponding analysis including multimers (**Figure S8**), although confirming the challenge of proline conformations, produced more variable results due to some hydrophobic residues poorly described due to location on multimer interfaces (i.e. exposed hydrophobic residues).

### AF2 prediction of residues in multimeric structures

The AF2 predictions are all single chains whereas most of the experimental structures are multimeric, either homomultimers or heteromultimers. We showed above that the correlation between RSA_AF_ and RSA_Exp_ values is substantially stronger for experimental monomers compared to multimers, and that the difference seems to be caused by residues that have low RSA_Exp_ but high RSA_AF_ (**Figure 2**). This makes intuitive sense as residues located in the interface between two chains may have lower solvent accessibility.

To test if the lower correlation between RSA_AF_ and RSA_Exp_ values observed in multimeric data pairs are caused by residues in chain interfaces, we identified such residues in the experimental structures and removed them from analysis. Chain interface residues were defined as residues whose atoms had a distance <3.5 Å to atoms in other chains. As can be seen in the left panels of **Figure 5,** the correlation between RSA_AF_ and RSA_Exp_ for interface residues is weak. The average RSA_AF_ for all identified interface residues in the multimer dataset (*n* = 2751) is 0.323 whereas the average RSA_Exp_ (using the per-AF2-averaged RSA_Exp_ values) is 0.135.

**Figure 5.**
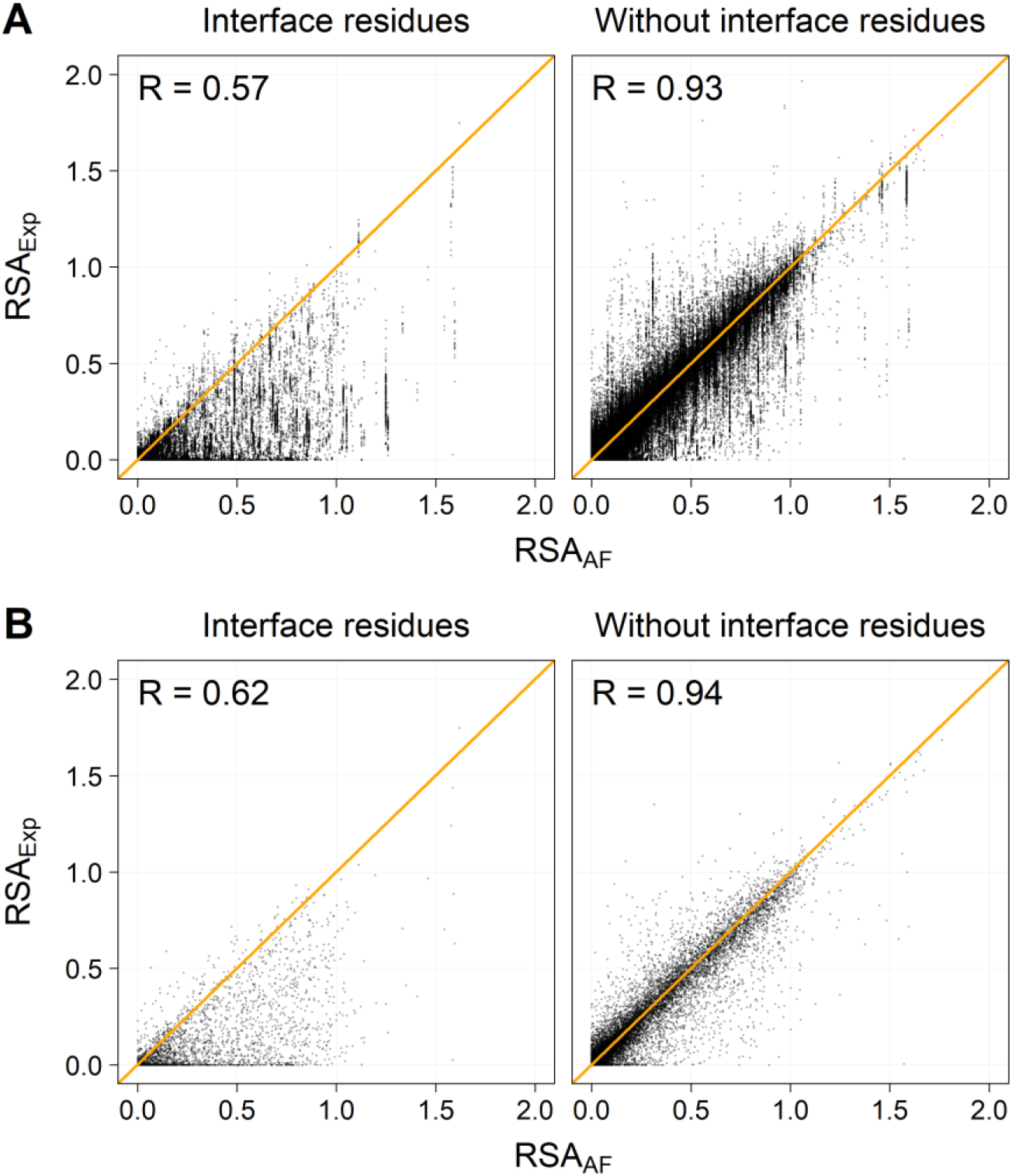
Experimental vs. AF2 RSA values in multimeric experimental structures. Correlation of RSA_AF_ and RSA_Exp_ for interface residues (left) and for non-interface residues (right). Orange lines represent the ideal where the RSA_AF_ are equal to RSA_Exp_. **(A)** Each dot represents a residue belonging to a data pair. **(B)** Each dot represents the average of experimental residues belonging to the same AlphaFold structure.

Thus, AF2 predictions on single chains highly overestimates the solvent accessibility of these residues. If interface residues are removed from the multimeric data pairs, the average RSA_AF_ of the remaining residues (*n* = 17,895) becomes 0.199 compared to an average RSA_Exp_ (using the per-AF2 averaged RSA_Exp_ values) of 0.190. This also results in a stronger correlation between RSA_AF_ and RSA_Exp_ that mimics the correlation of monomeric data pairs (compare right panels of **Figure 5** to upper panels of **Figure 2**). In summary, because the AF2 predictions are based on single protein chains, it overestimates the solvent accessibility of the ~10% of residues in the dataset that are in close contact (by the above definition) to other protein chains, and this should always be accounted for in predictions.

### Variation in RSA values among experimental structures of the same protein chain

Some AF2-structures are matched to many experimental structures (**Figure S3**). The different experimental structures representing one particular protein chain (and therefore one AF2-structure) can be compared to each other to determine the amount of variation in RSA values that exists for each residue among experimental structures. To evaluate the variation within the experimental dataset used to assess the prediction accuracy of AF2, we calculated the per-residue spread in RSA_Exp_ and compared it to the per-residue MAE for AF2-structures matched to five or more monomeric experimental structures (*n* = 10).

An example is shown in **Figure S9** for peptidyl prolyl *cis/trans* isomerase A (UniProt ID P62937) for which there are 52 monomeric experimental structures in the dataset. **Figure S9A** shows the SD of RSA_Exp_ (SD_Exp_) for each residue in the chain. Although the experimental structures are all monomers, and variation caused by chain interactions are therefore not relevant, some residues have very different RSA values in the 52 structures while others only show little RSA variation. For comparison, **Figure S9B** shows the absolute errors between RSA_AF_ and RSA_Exp_ for each residue along the protein chain.

To better understand how experimental variation affects the performance benchmark, **Figure 6A** shows correlation between the per-residue variation in RSA_Exp_ (measured as SD_Abs_) and the per-residue MAE for the 2636 compared residues. The outliers in **Figure 6A** can be interpreted in terms of confidence in the RSA_AF_ prediction. Residues for which the RSA is similar among the experimental structures (low SD_Abs_) but different from the AF2-predicted RSA (high MAE) can be interpreted as residues that are unlikely to be predicted accurately. In contrast, residues for which there is a high variation in RSA among the experimental structures (high SD_Abs_) but a low MAE when compared to the AF2-prediction, indicate residues for which some experimental structures may be problematic.

**Figure 6.**
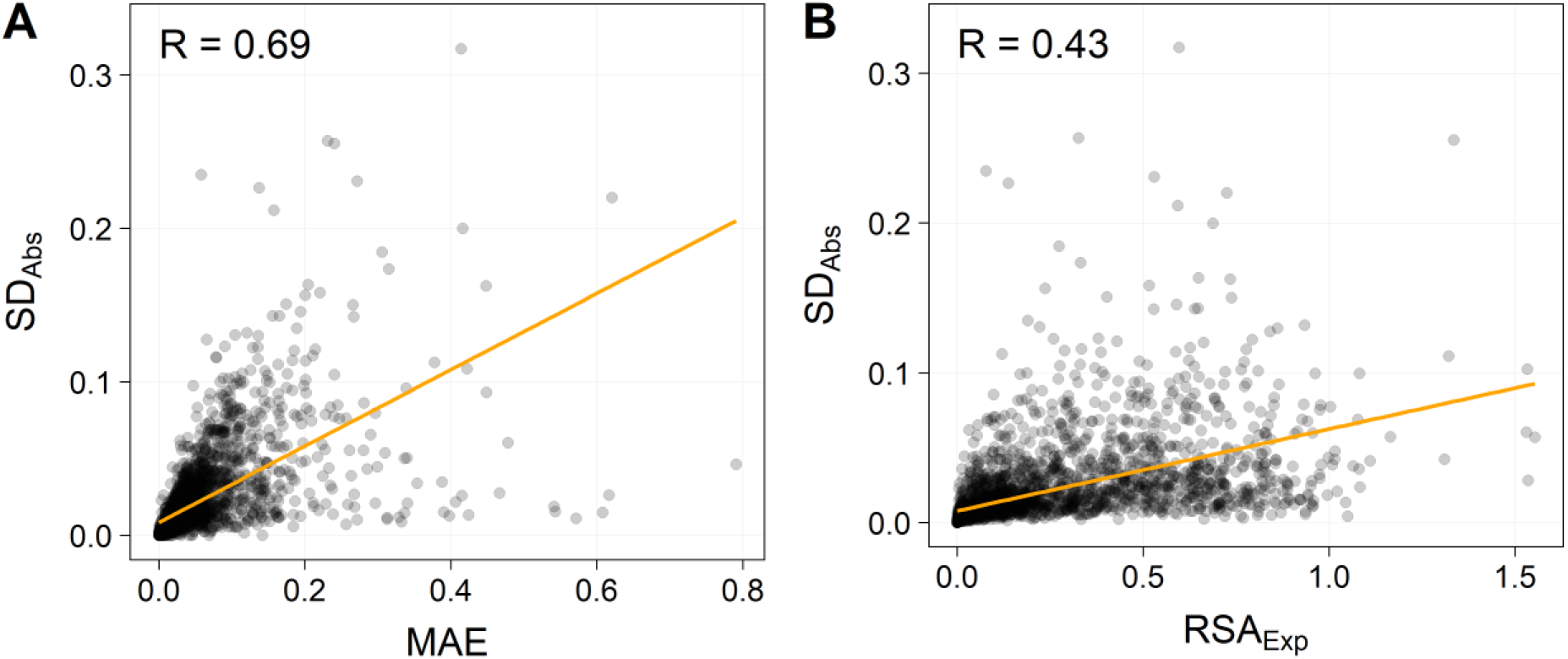
Internal variation among experimental structures. Each point indicates one residue in the monomeric data pairs for which there are 5 or more experimental structures per AlphaFold structure **A)** Correlation of SD_Abs_ and MAE, **B)** correlation between SD_Abs_ and RSA_Exp_. Linear regressions and correlation coefficients (R) are shown.

Furthermore, **Figure 6B** shows that the variation in RSA among experimental structures depends to some extent on RSA_Exp_ with more surface exposed residues exhibiting more variation. This structural heterogeneity in experimental structures not only puts a limit on the accuracy expected from a prediction method but also emphasizes the need to consider the experimental structure quality and heterogeneity in an assessment, e.g. by sensitivity analysis or precision estimates using multiple experimental structures in the study.

### Concluding remarks

We analyzed the performance of AF2 applied to human proteins, using the local residue’s relative solvent exposure as a “natural” metric of its conformation. We carefully curated data sets by sequence overlap to avoid biases from incomplete or erroneous structures and explored the dependence of AF2 performance on monomer/multimer status (important), presence of cofactors and ligands, experimental resolution, exposure, and amino acid type. AF2 performed excellently once comparing specifically to monomer proteins. However, notable challenges persist relating especially to proline and exposed residues. We identify larger errors for lower-confidence scores (pLDDT) and exposed residues on average, which also correlates with polar residues (Asp, Glu, Asn e.g.) being substantially less well described than hydrophobic residues. We provide expected errors divided into amino acid type, exposure, and pLDDT value, and emphasize the structural heterogeneity of experimental data.

An important point of caution is that in the application to real proteomes, the predictions of AF2 are likely worse due to training on usually unmodified or specifically modified proteins from the PDB, whereas many eukaryotic protein chains are heavily modified (truncated, glycosylated, etc.) sometimes in very diverse ways in their physiologically relevant forms (Bagdonas *et al*., 2021). To this, we may also add the fact that the protein structures used for training and assessment represent crystal state in vitro conditions not necessarily applicable to the pH, temperature, and macromolecular cellular environment where the protein is located. Thus, we are still far from in vivo predictability of protein conformational ensembles - but we are possibly close to understand in vitro protein conformational ensembles, as this study has hopefully helped to document.

## Supporting information

Table S1 performance data

## Acknowledgement

The Danish Council for Independent Research (Grant case: 8022-00041B) is gratefully acknowledged for supporting this work.

## Supporting Information

Table S1 (csv file) contains data and performance metrics. The supporting information pdf file contains additional tables and figures with information relevant to analysis (Table S2, Table S3, and Figures S1-S9).

## Data availability

All the data used and produced in this work are described in the Supporting Information. All data analyses were performed with Python v.3.8 or R v.4.0.5 and the code is available at https://github.com/ktbaek/AlphaFold.

