## Supporting information figures and tables for "Assessment of AlphaFold2 residue conformations for human proteins"

*Kasper P. Kepp: 0000-0002-6754-7348*

*Kristoffer T. Bæk: 0000-0003-3243-3962*

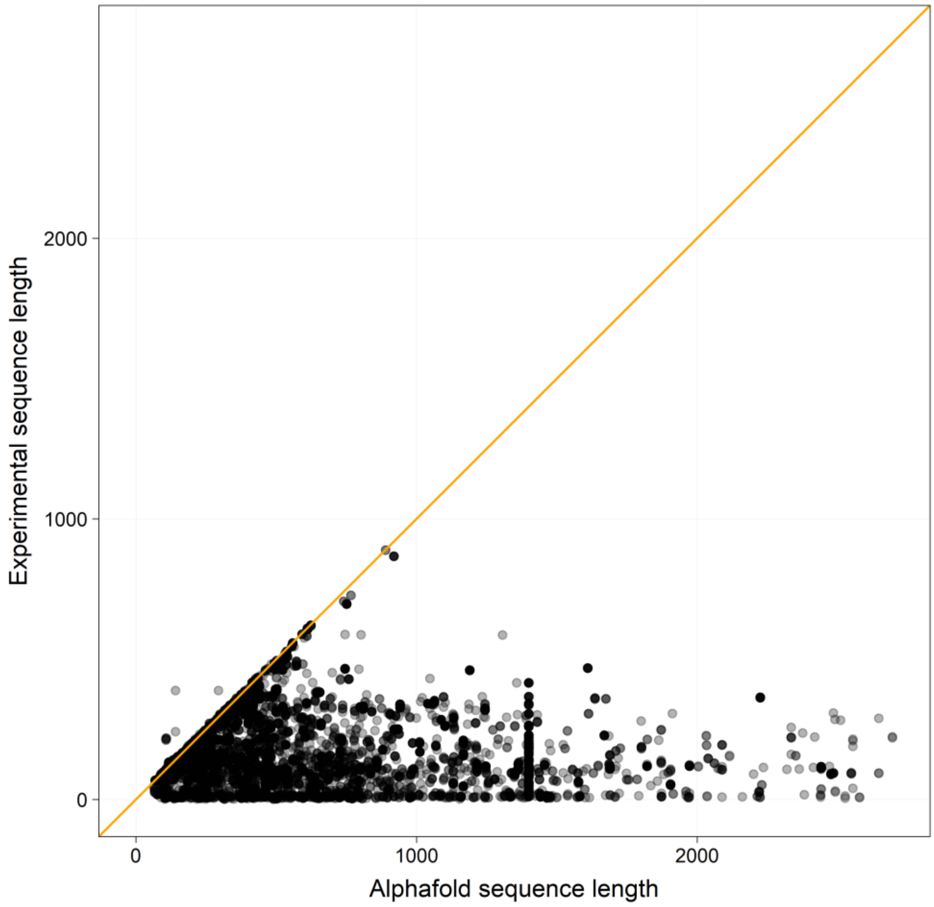


**Figure S1. AlphaFold structure sequence length vs experimental structure sequence length.** Each data point represents a data pair matched by UniProt number. Only experimental structures with no internal gaps, insertions or mutations are included.

Figure S2 (1)

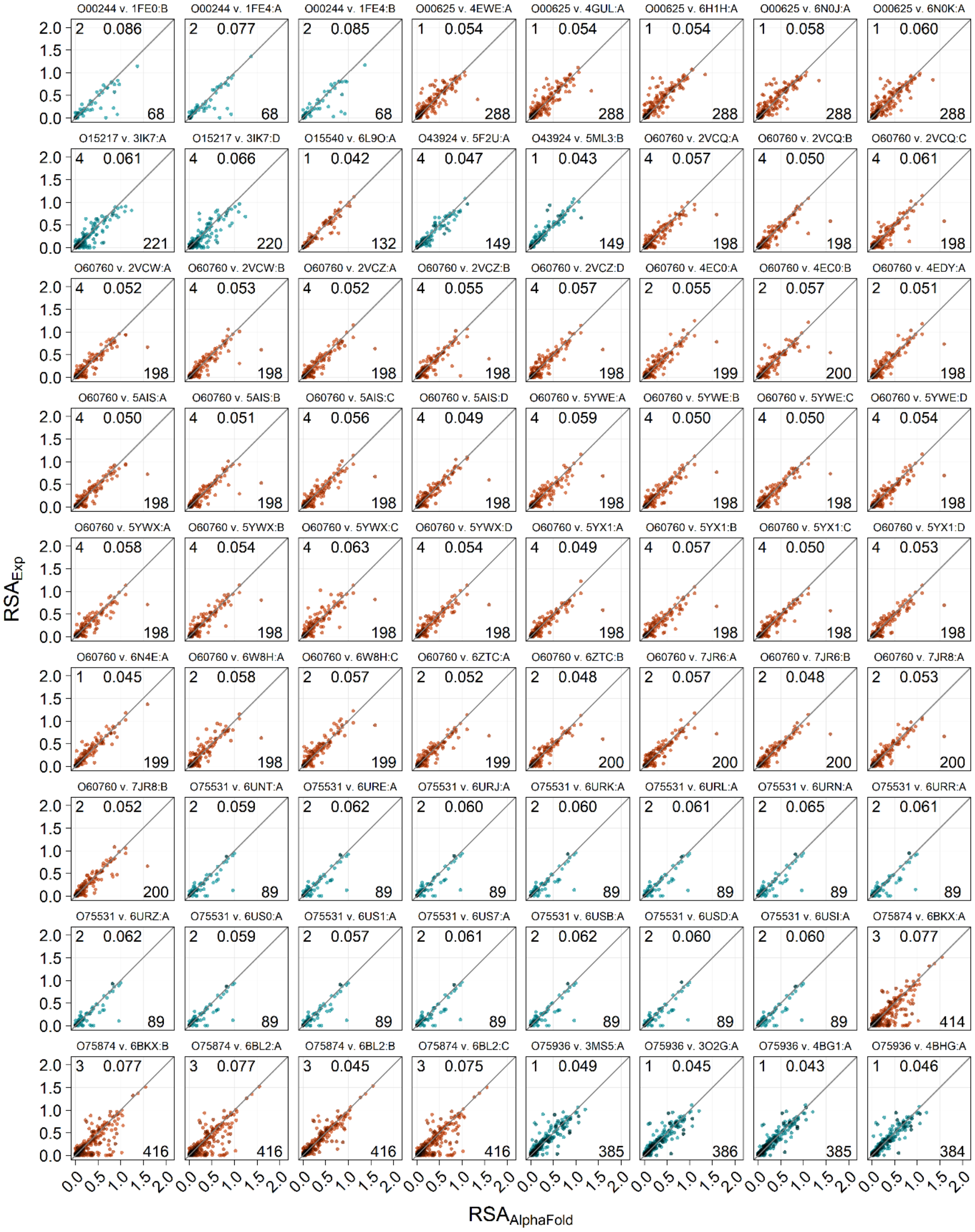
O00244 v. 1FE0:B
O00244 v. 1FE4:A
O00244 v. 1FE4:B
O00625 v. 4EWE:A
O00625 v. 4GUL:A
O00625 v. 6H1H:A
O00625 v. 6N0J:A
O00625 v. 6N0K:A
O15217 v. 3IK7:A
O15217 v. 3IK7:D
O15540 v. 6L9O:A
O43924 v. 5F2U:A
O43924 v. 5ML3:B
O60760 v. 2VCQ:A
O60760 v. 2VCQ:B
O60760 v. 2VCQ:C
O60760 v. 2VCW:A
O60760 v. 2VCW:B
O60760 v. 2VCZ:A
O60760 v. 2VCZ:B
O60760 v. 2VCZ:D
O60760 v. 4EC0:A
O60760 v. 4EC0:B
O60760 v. 4EDY:A
O60760 v. 5AIS:A
O60760 v. 5AIS:B
O60760 v. 5AIS:C
O60760 v. 5AIS:D
O60760 v. 5YWE:A
O60760 v. 5YWE:B
O60760 v. 5YWE:C
O60760 v. 5YWE:D
O60760 v. 5YWX:A
O60760 v. 5YWX:B
O60760 v. 5YWX:C
O60760 v. 5YWX:D
O60760 v. 5YX1:A
O60760 v. 5YX1:B
O60760 v. 5YX1:C
O60760 v. 5YX1:D
O60760 v. 6N4E:A
O60760 v. 6W8H:A
O60760 v. 6W8H:C
O60760 v. 6ZTC:A
O60760 v. 6ZTC:B
O60760 v. 7JR6:A
O60760 v. 7JR6:B
O60760 v. 7JR8:A
O60760 v. 7JR8:B
O75531 v. 6UNT:A
O75531 v. 6URE:A
O75531 v. 6URJ:A
O75531 v. 6URK:A
O75531 v. 6URL:A
O75531 v. 6URN:A
O75531 v. 6URR:A
O75531 v. 6URZ:A
O75531 v. 6US0:A
O75531 v. 6US1:A
O75531 v. 6US7:A
O75531 v. 6USB:A
O75531 v. 6USD:A
O75531 v. 6USI:A
O75874 v. 6BKX:A
O75874 v. 6BKX:B
O75874 v. 6BL2:A
O75874 v. 6BL2:B
O75874 v. 6BL2:C
O75936 v. 3MS5:A
O75936 v. 3O2G:A
O75936 v. 4BG1:A
O75936 v. 4BHG:A
RSAExp
RSAAlphaFold

Figure S2, continued (2)

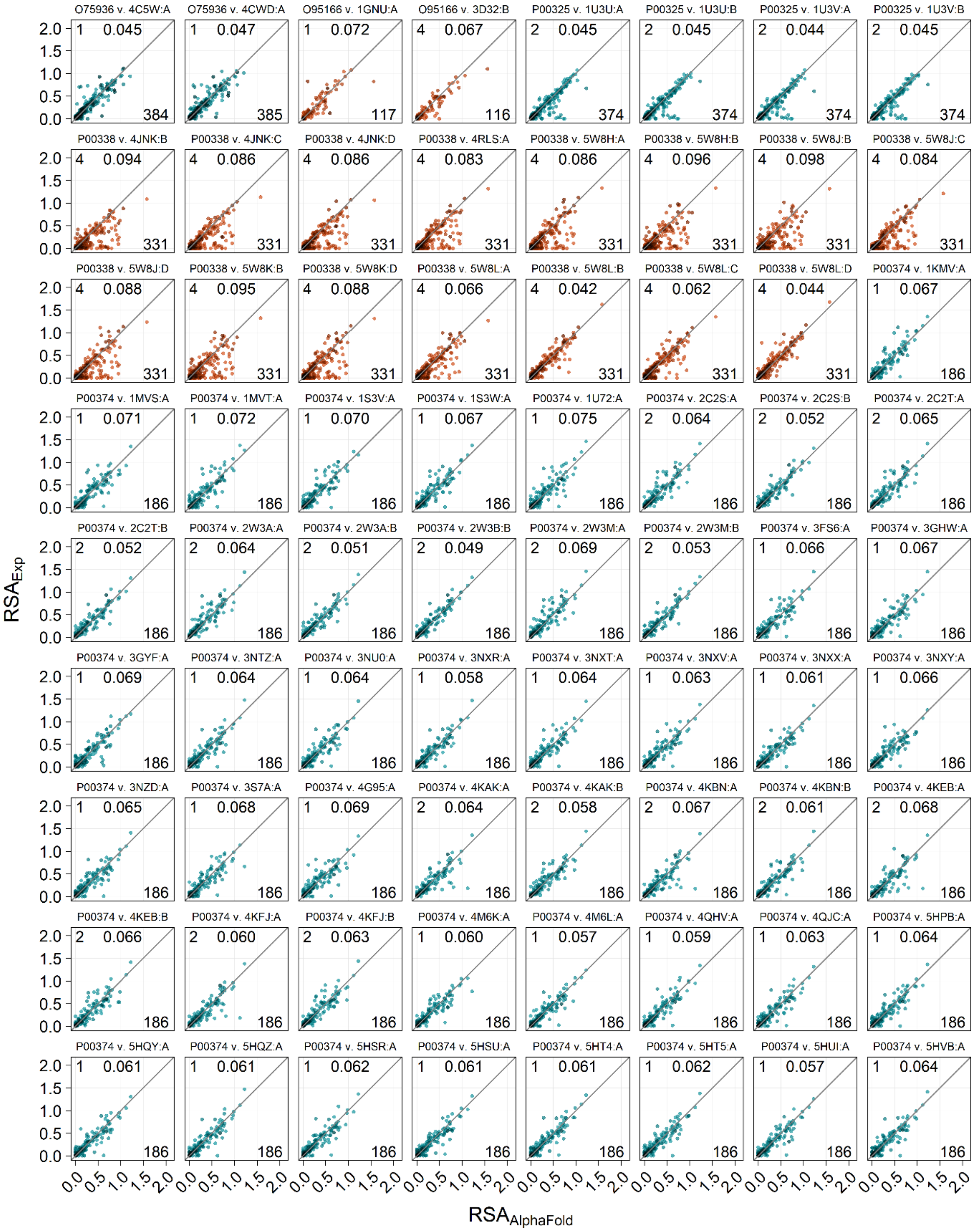

Figure S2, continued (3)

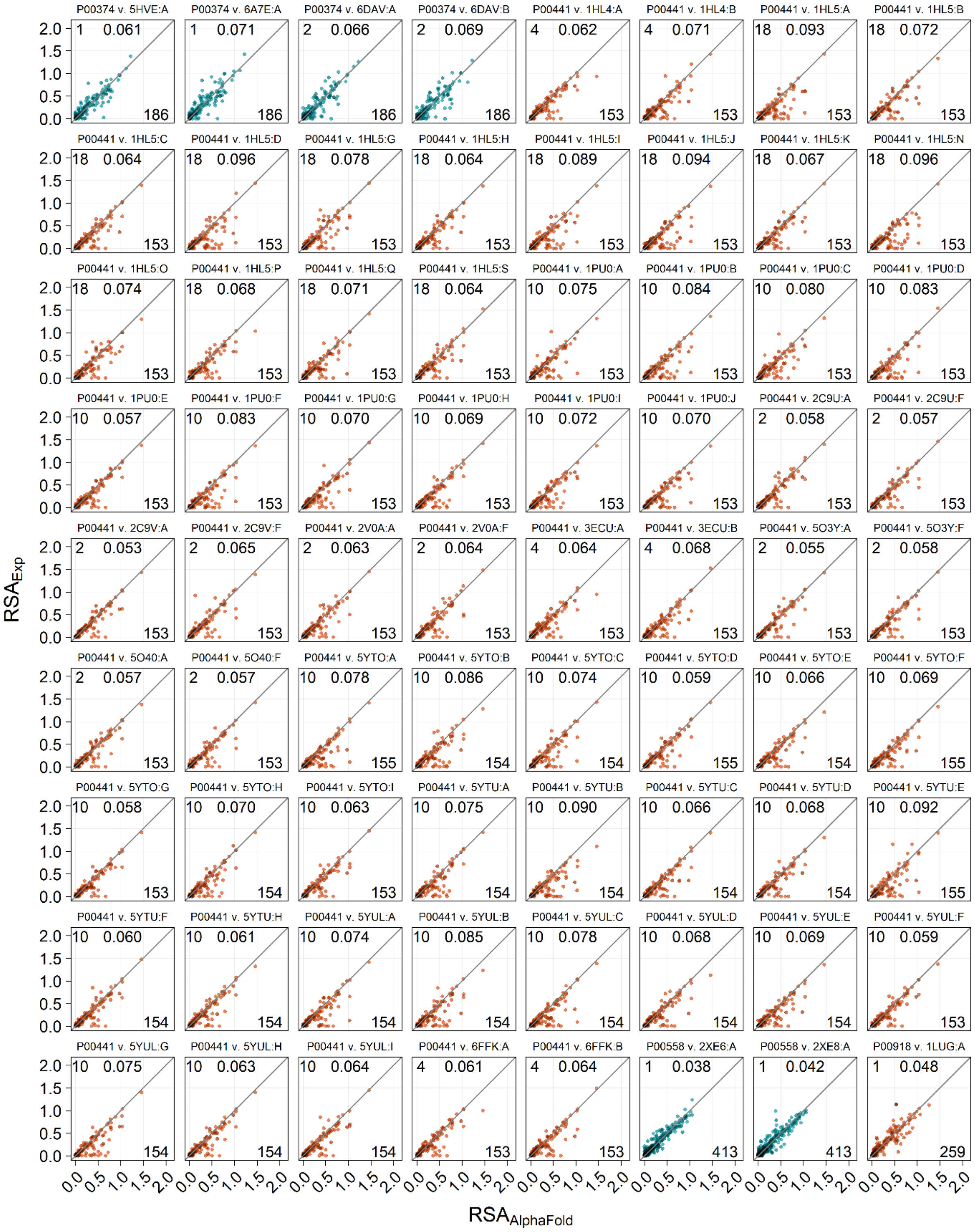

Figure S2, continued (4)

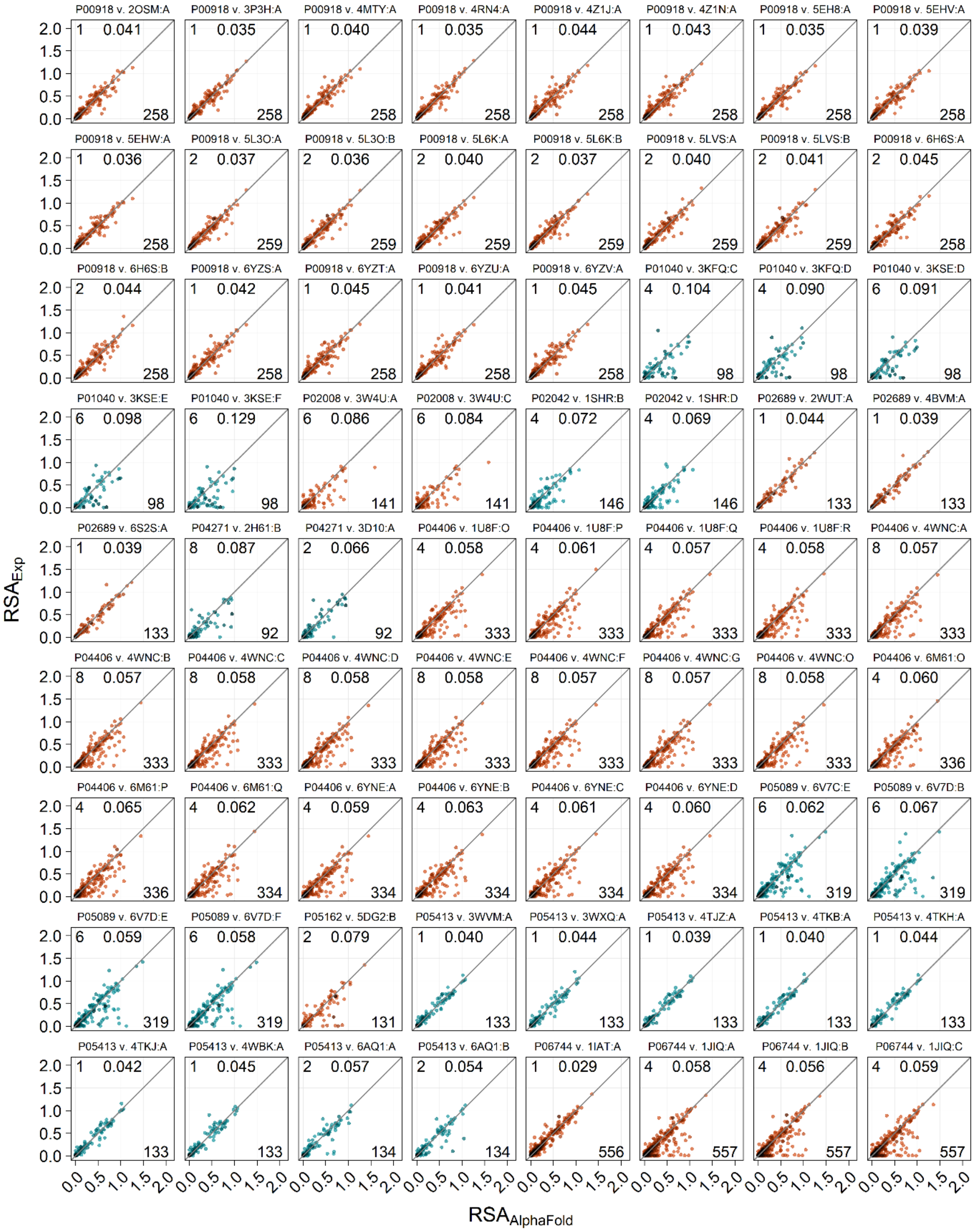

Figure S2, continued (5)

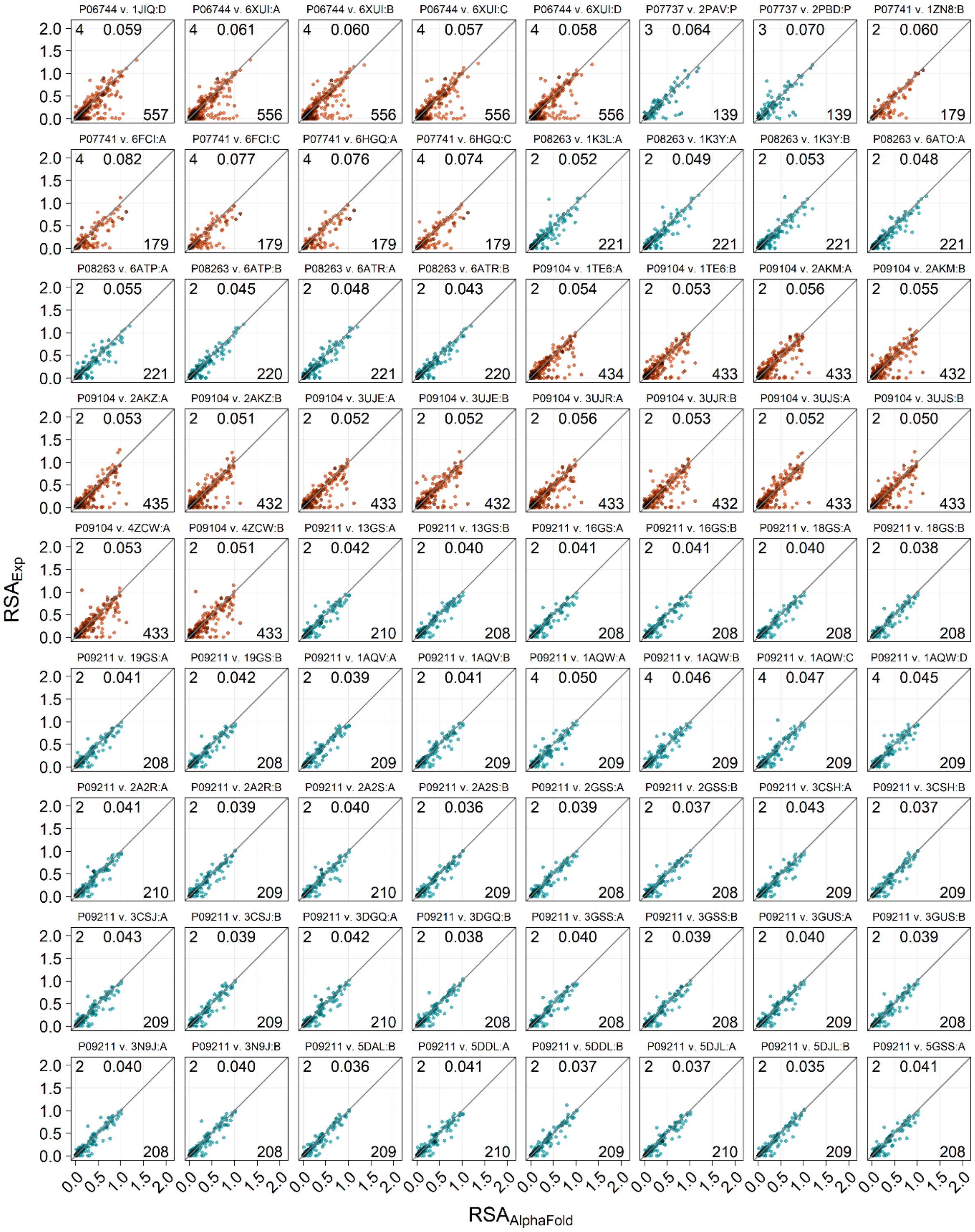

Figure S2, continued (6)

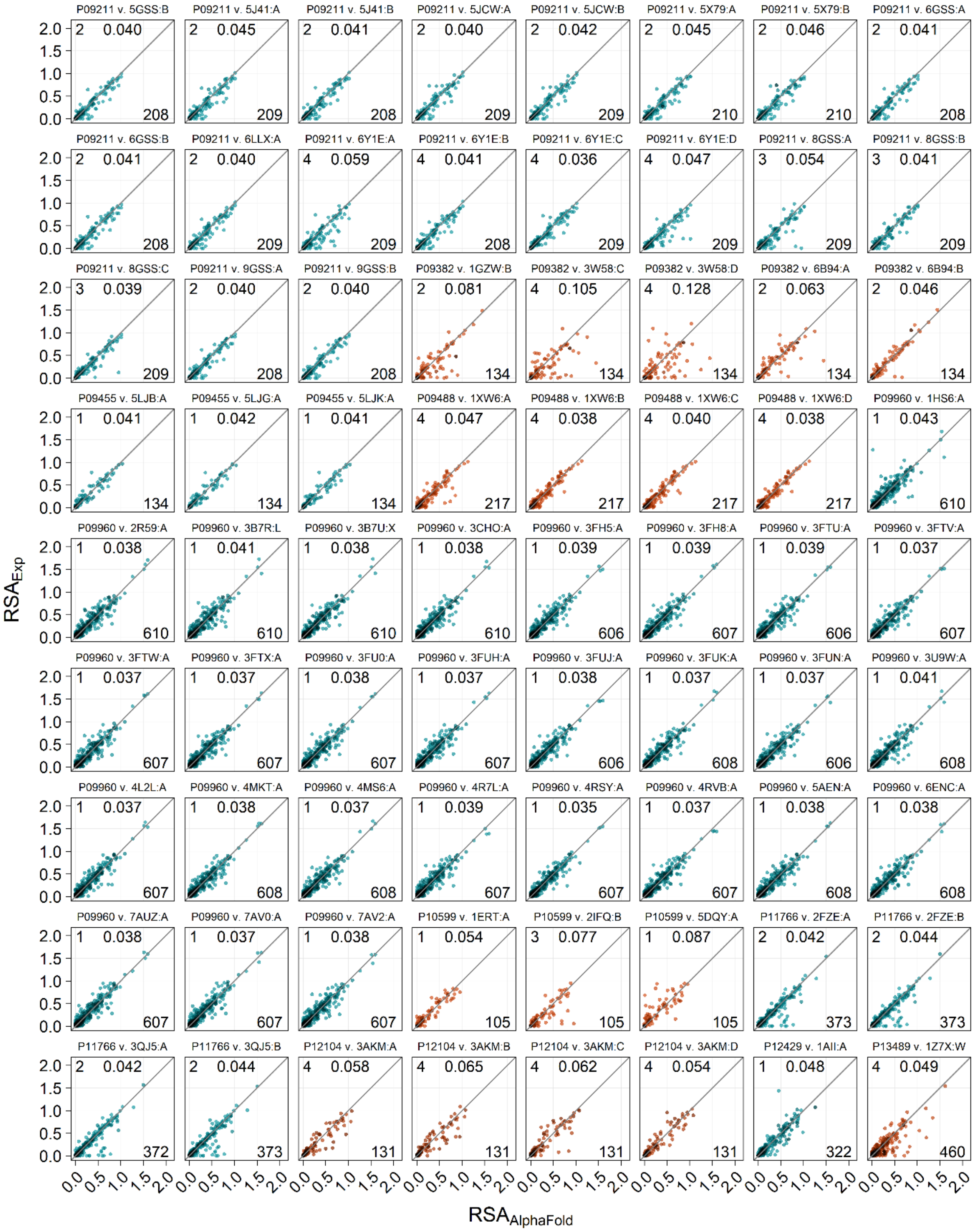

Figure S2, continued (7)

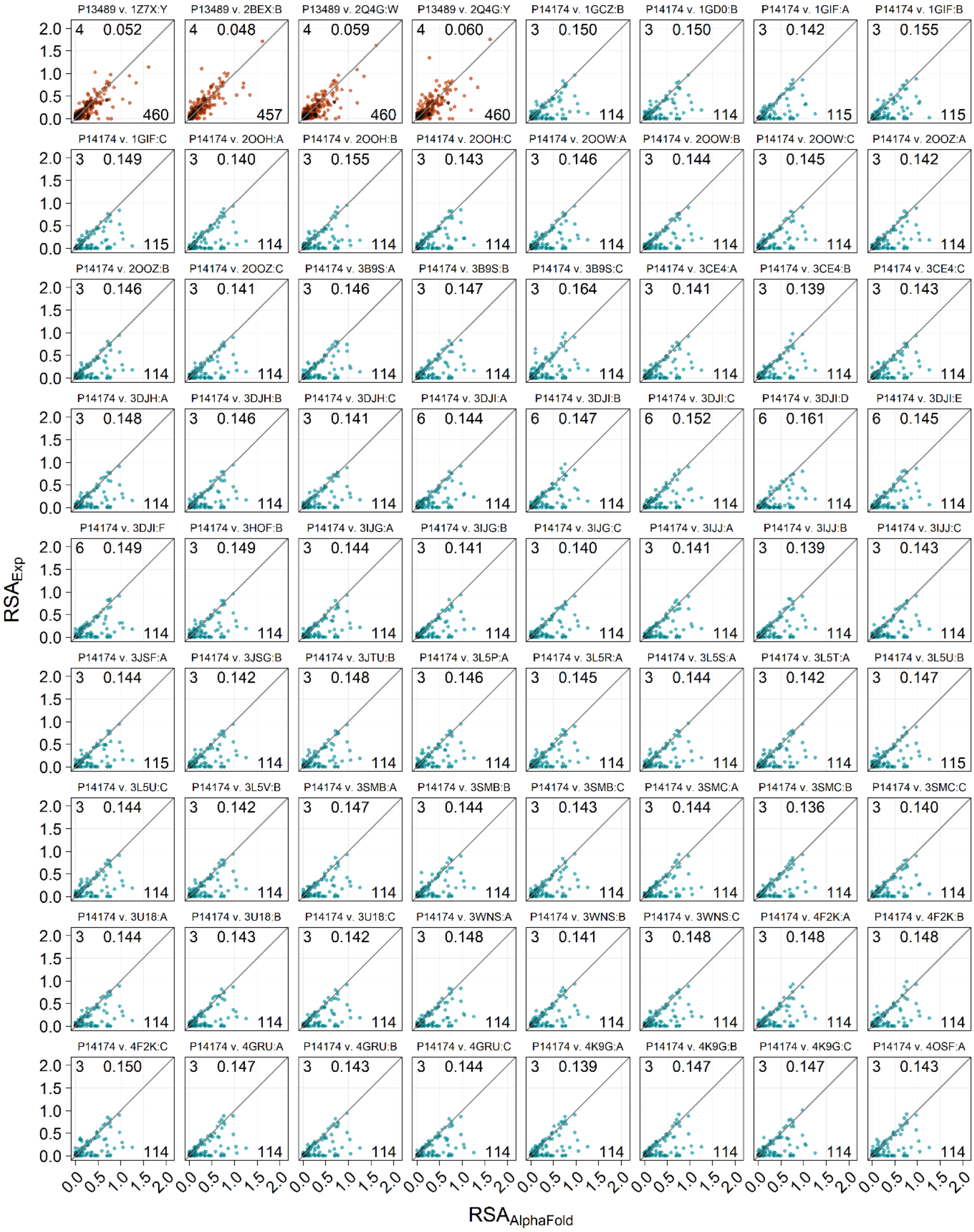

Figure S2, continued (8)

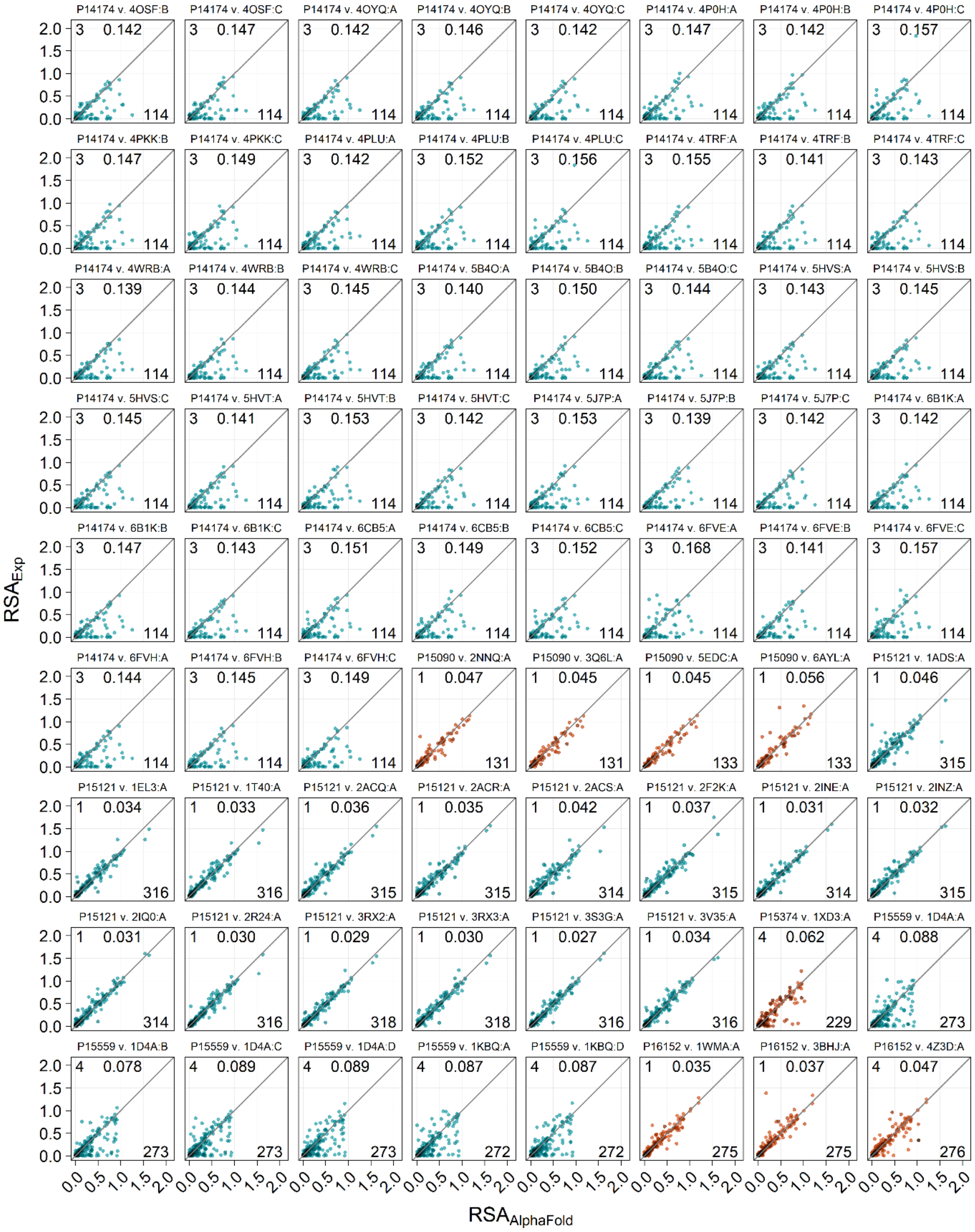

Figure S2, continued (9)

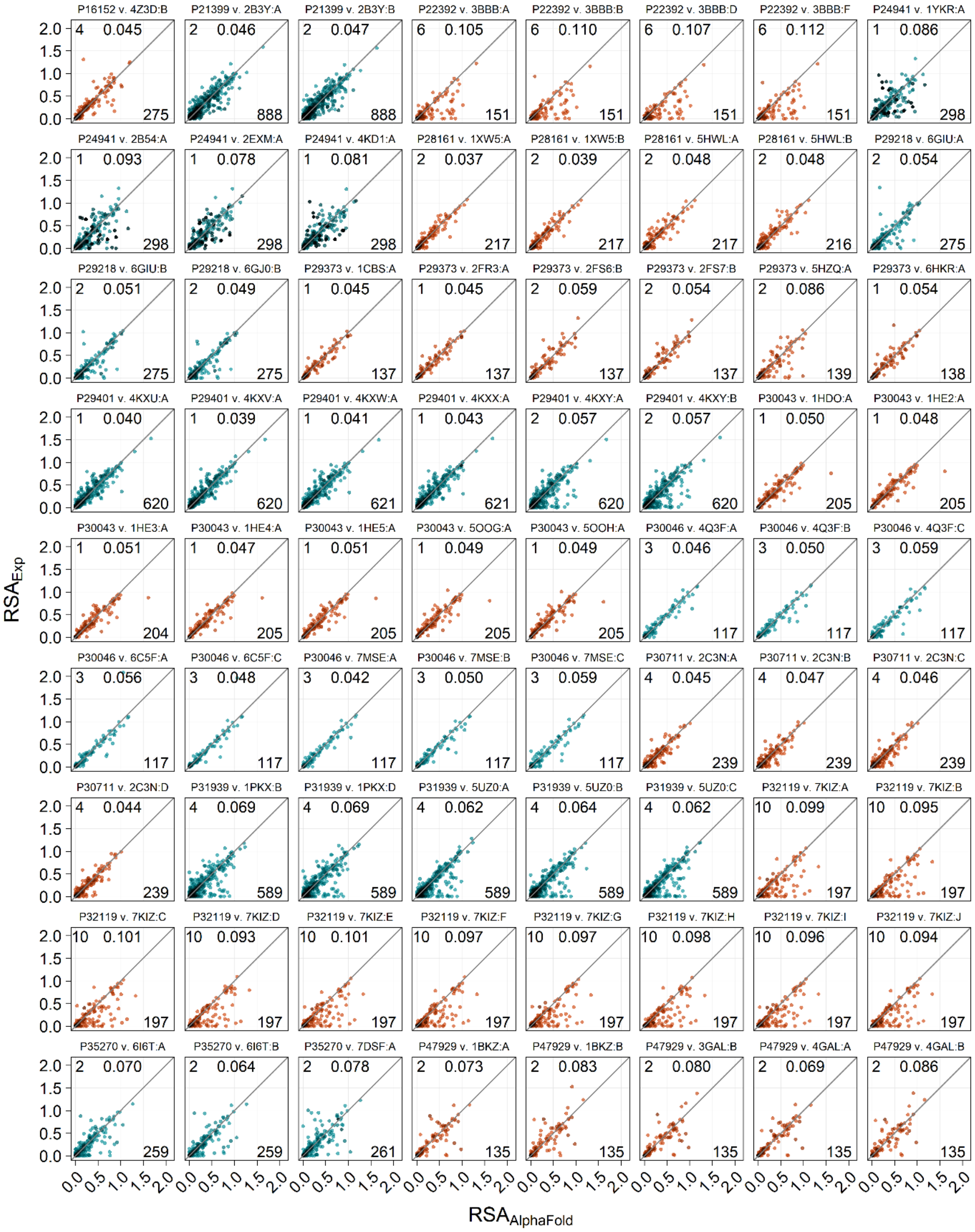

Figure S2, continued (10)

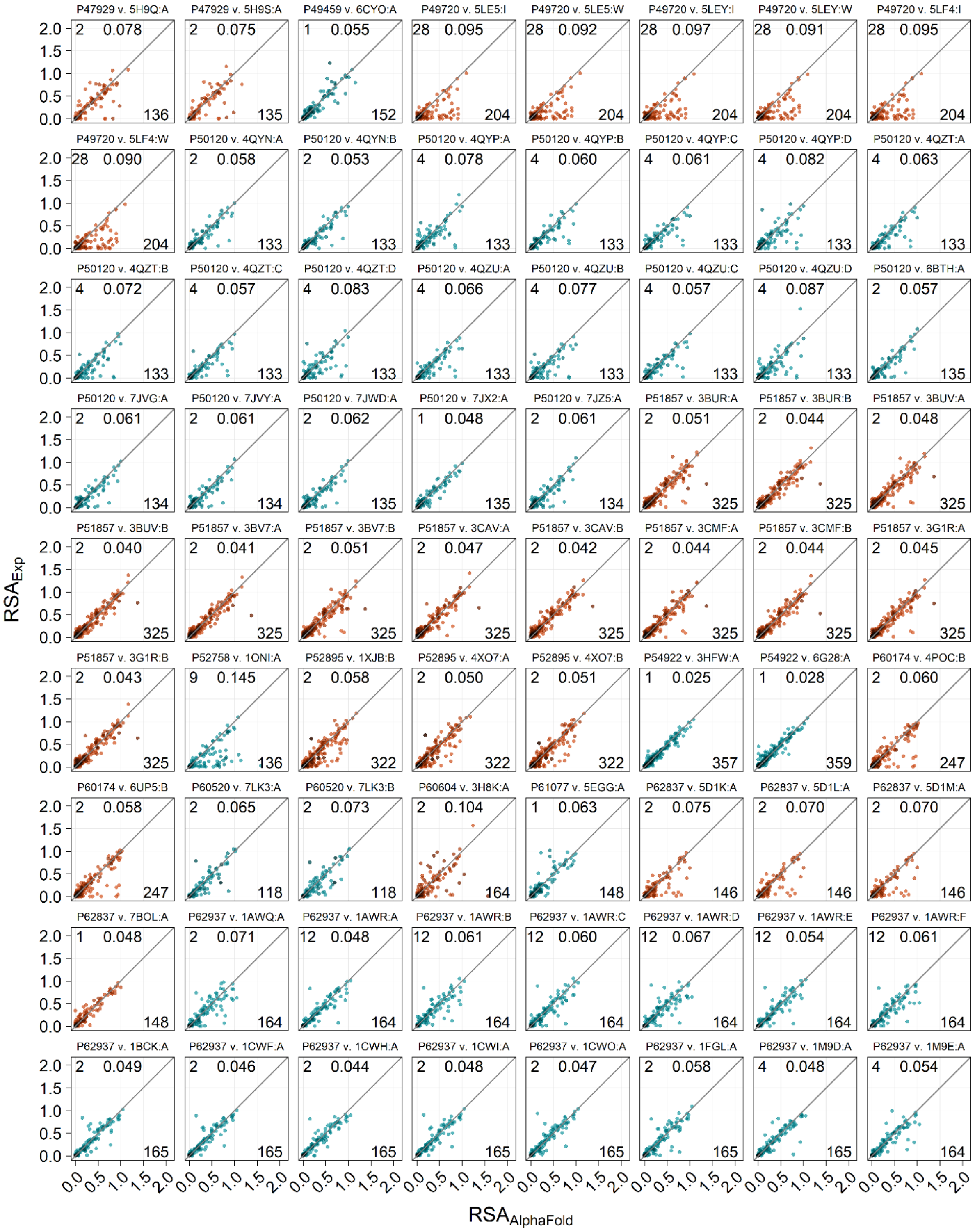

Figure S2, continued (11)

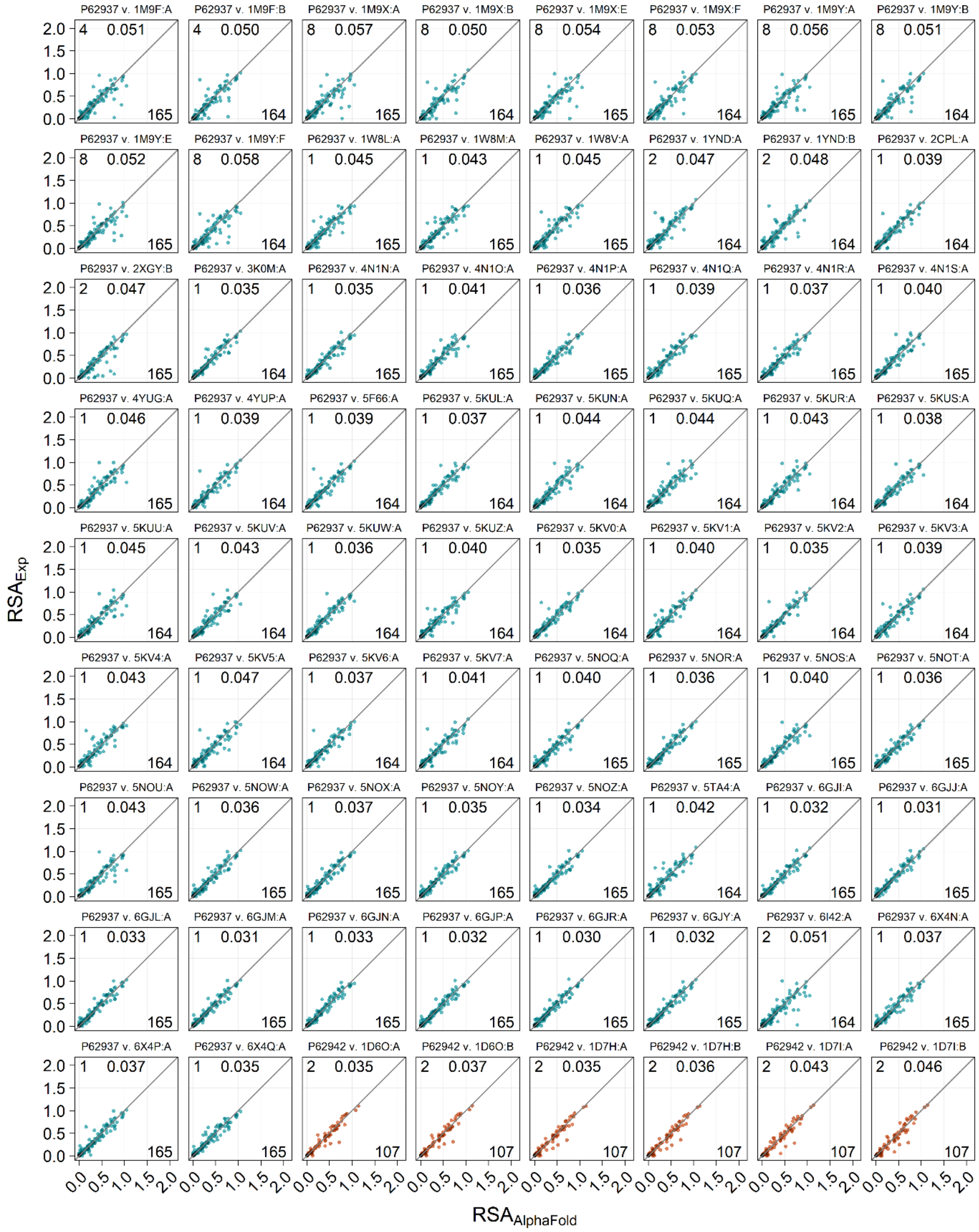

Figure S2, continued (12)

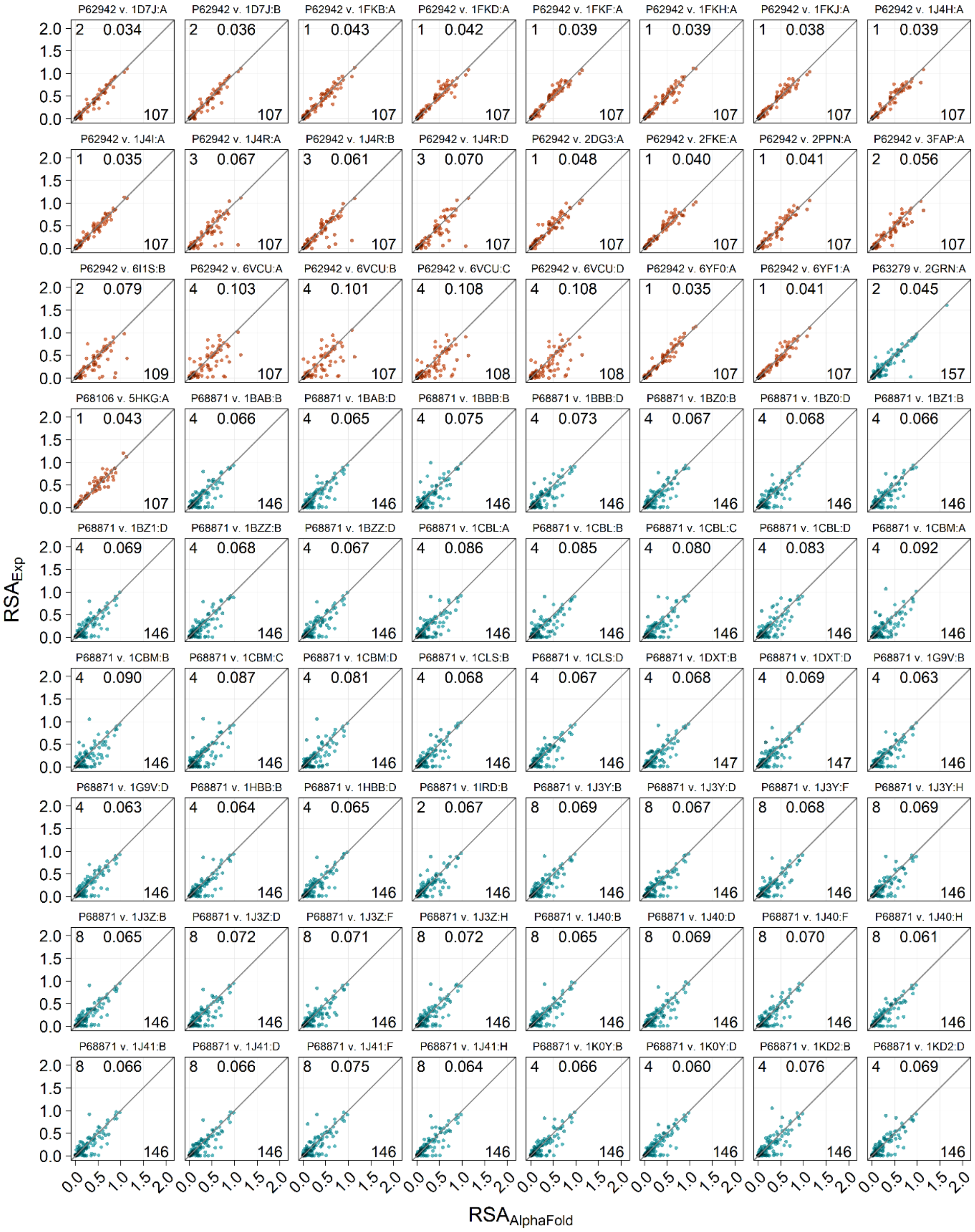

Figure S2, continued (13)

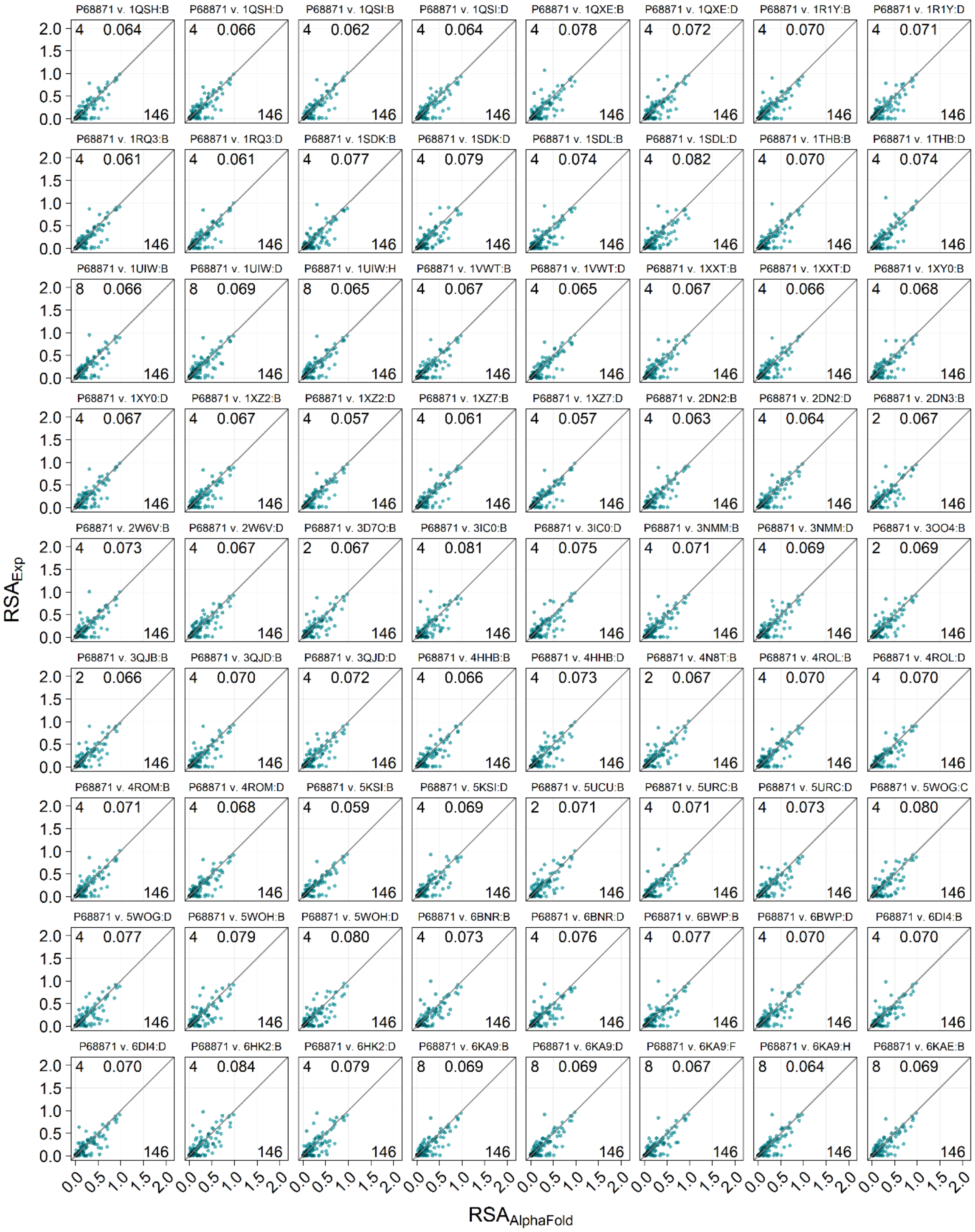

Figure S2, continued (14)

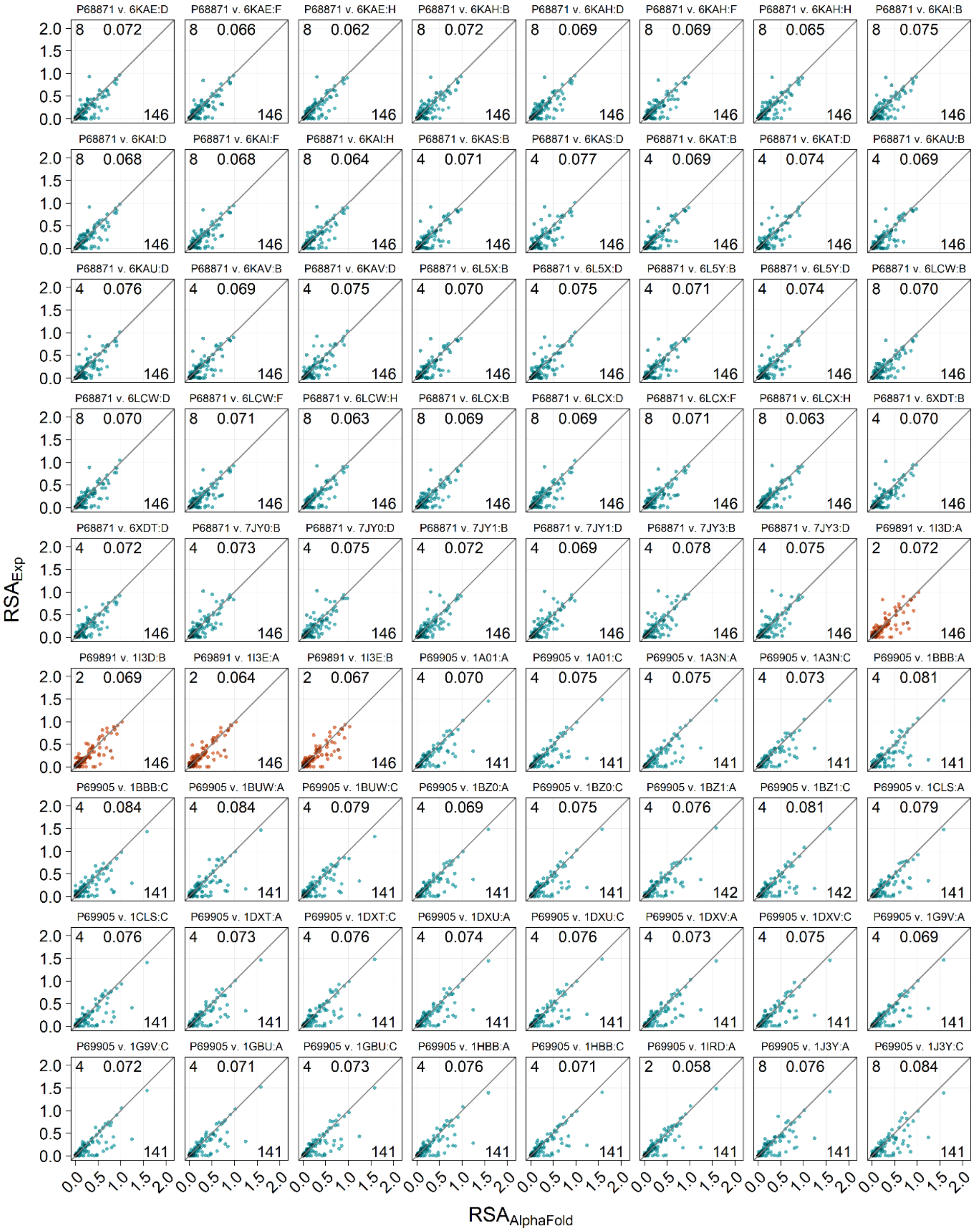

Figure S2, continued (15)

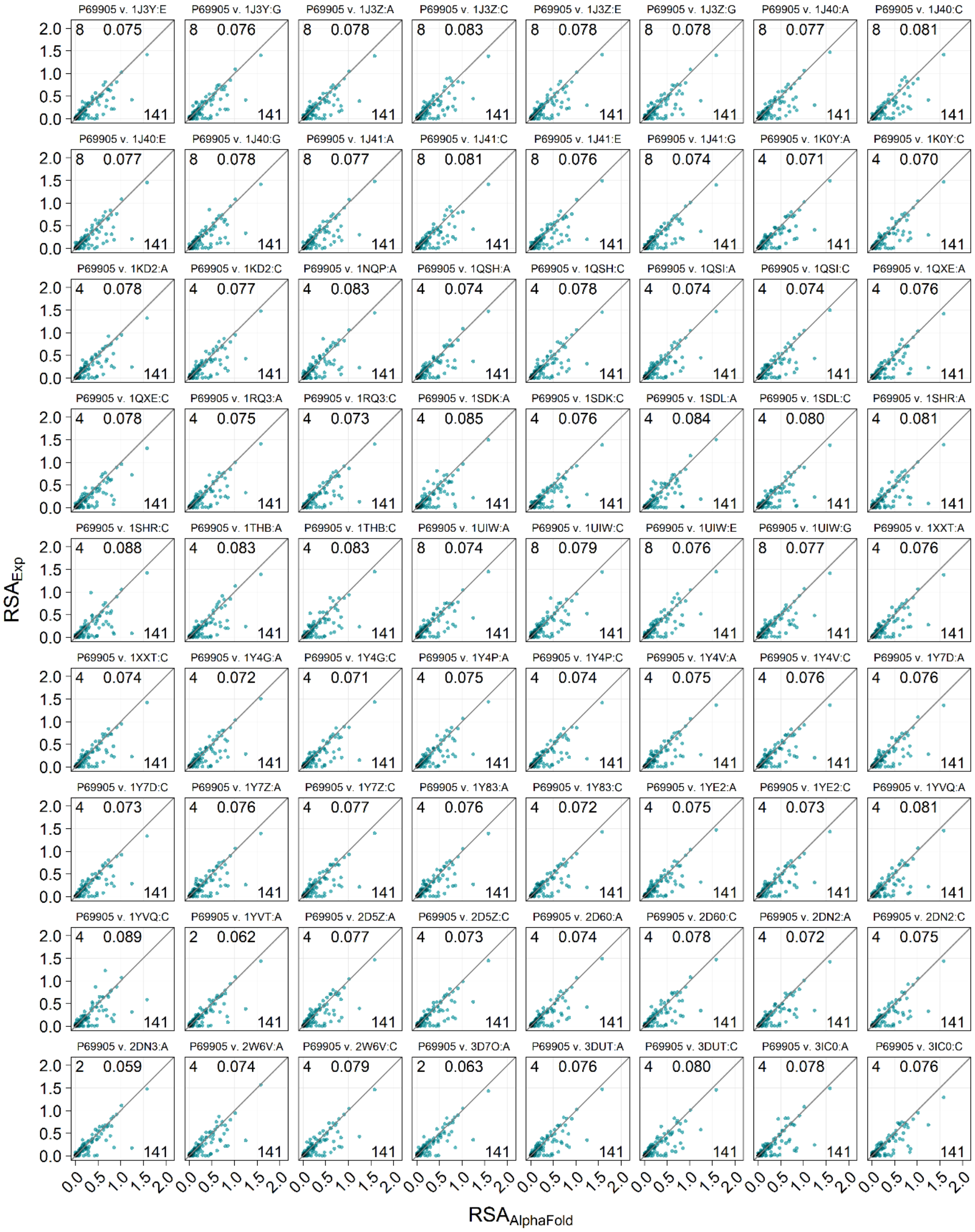

Figure S2, continued (16)

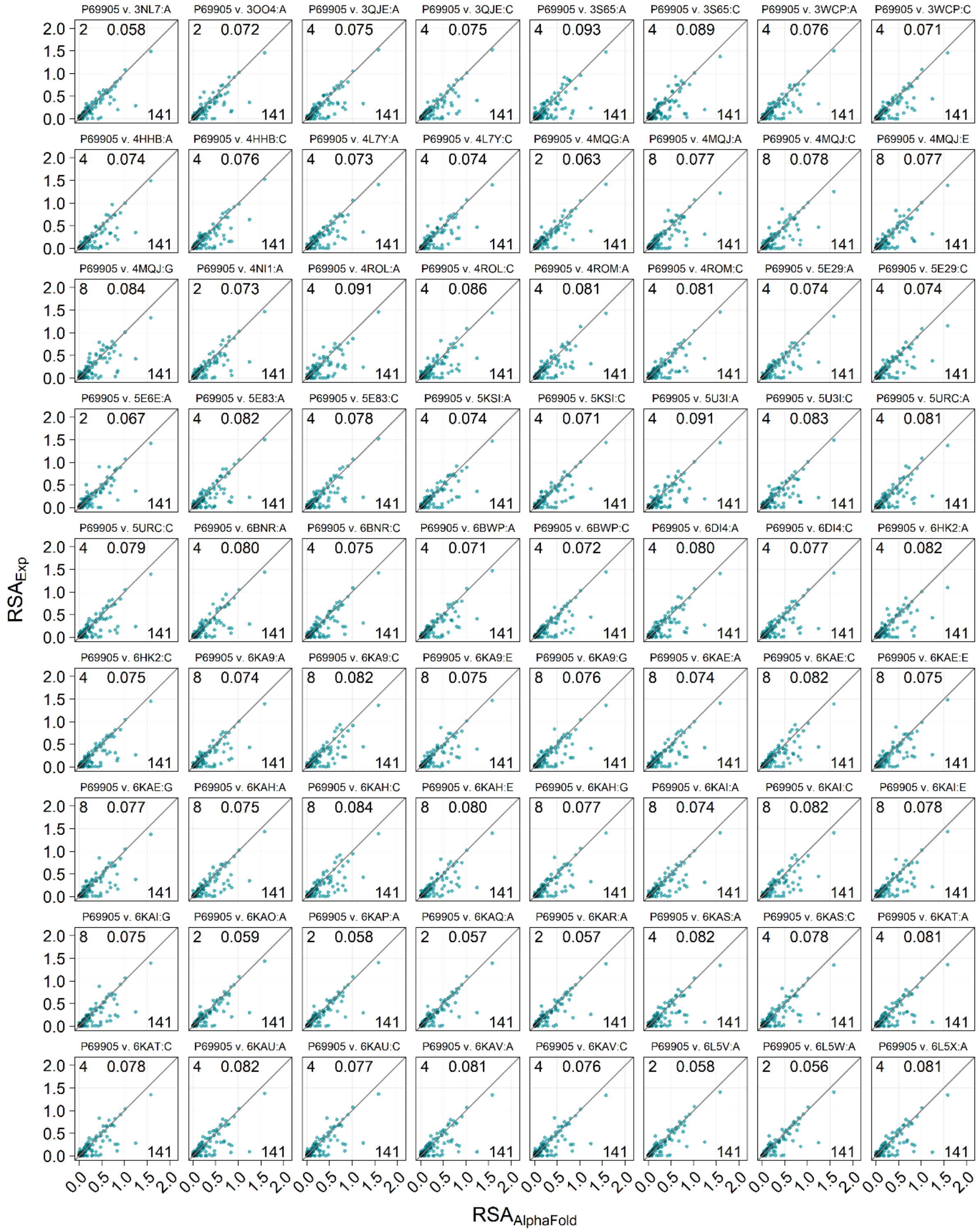

Figure S2, continued (17)

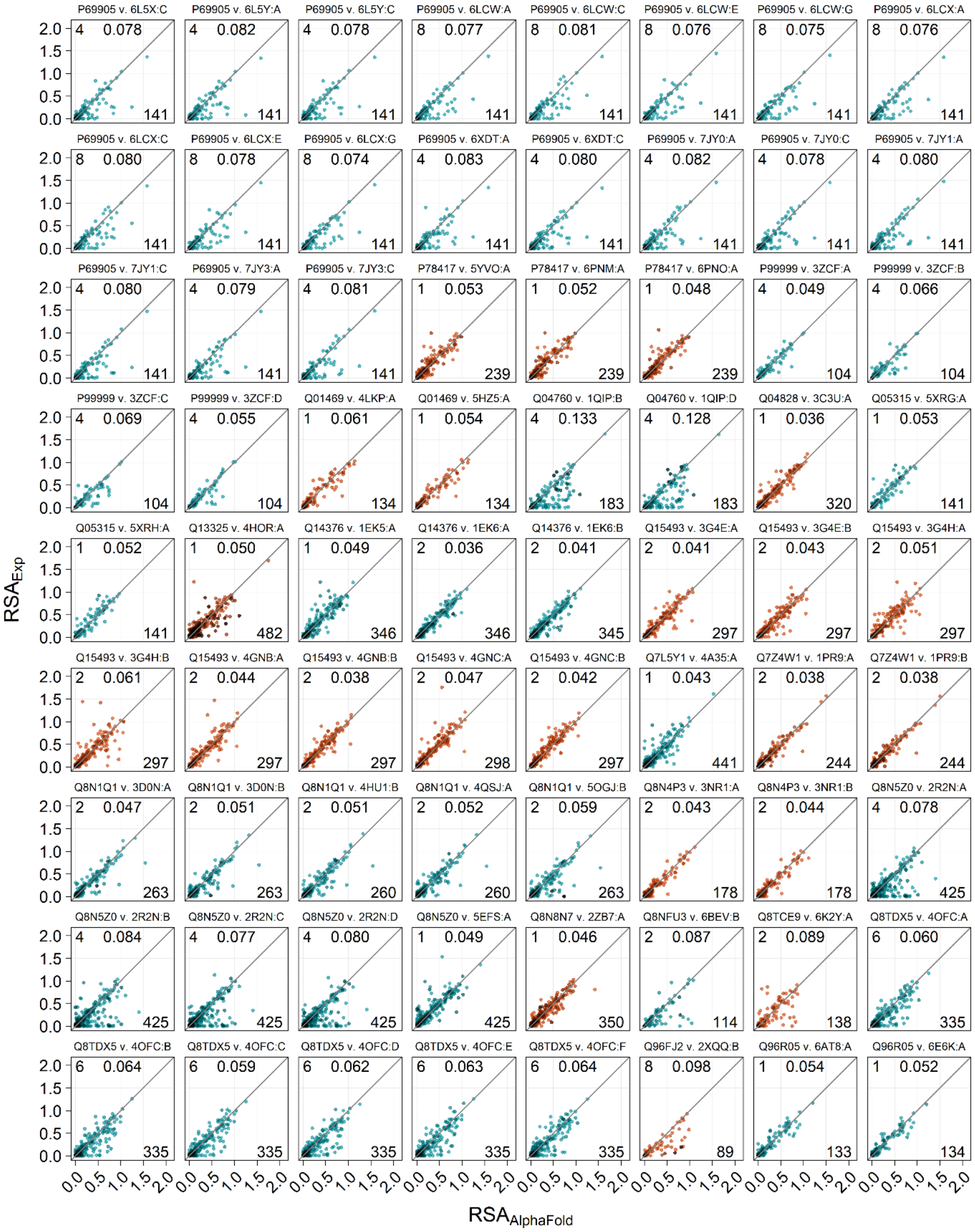

Figure S2, continued (18)

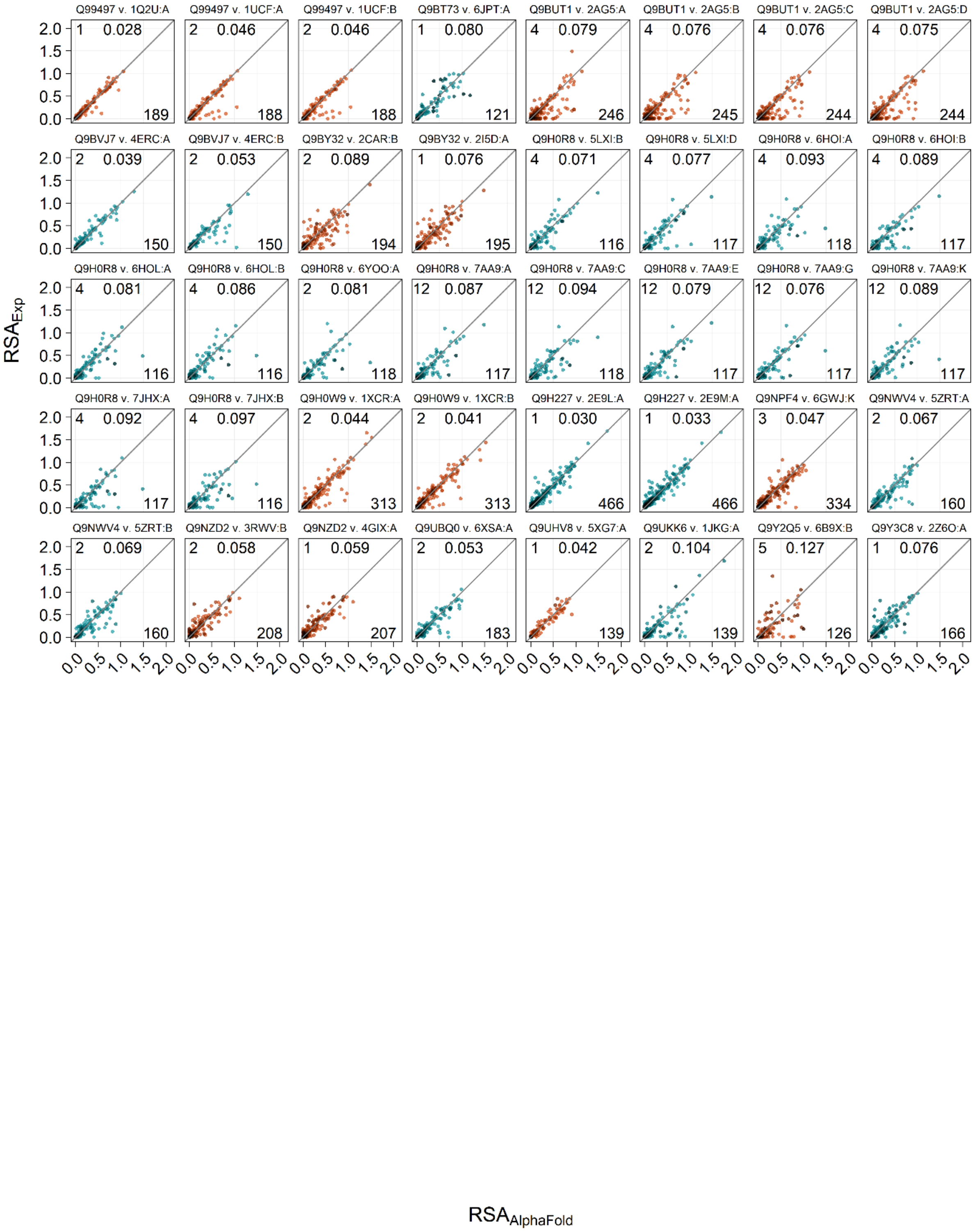

**Figure S2: Experimental RSA values vs AlphaFold RSA values.** Each data point represents a residue belonging to a data pair. Each panel shows one structure pair. The color of the points alternate with each unique AlphaFold structure for a better overview. Each point is furthermore color coded according to the pLDDT value: darker shades show lower pLDDT value i.e less confidence. The value in the upper left corner indicates the number of chains in the experimental structure, the value in the upper right corner indicates the MAE, and the value in the lower right corner indicates the number of matched residues.

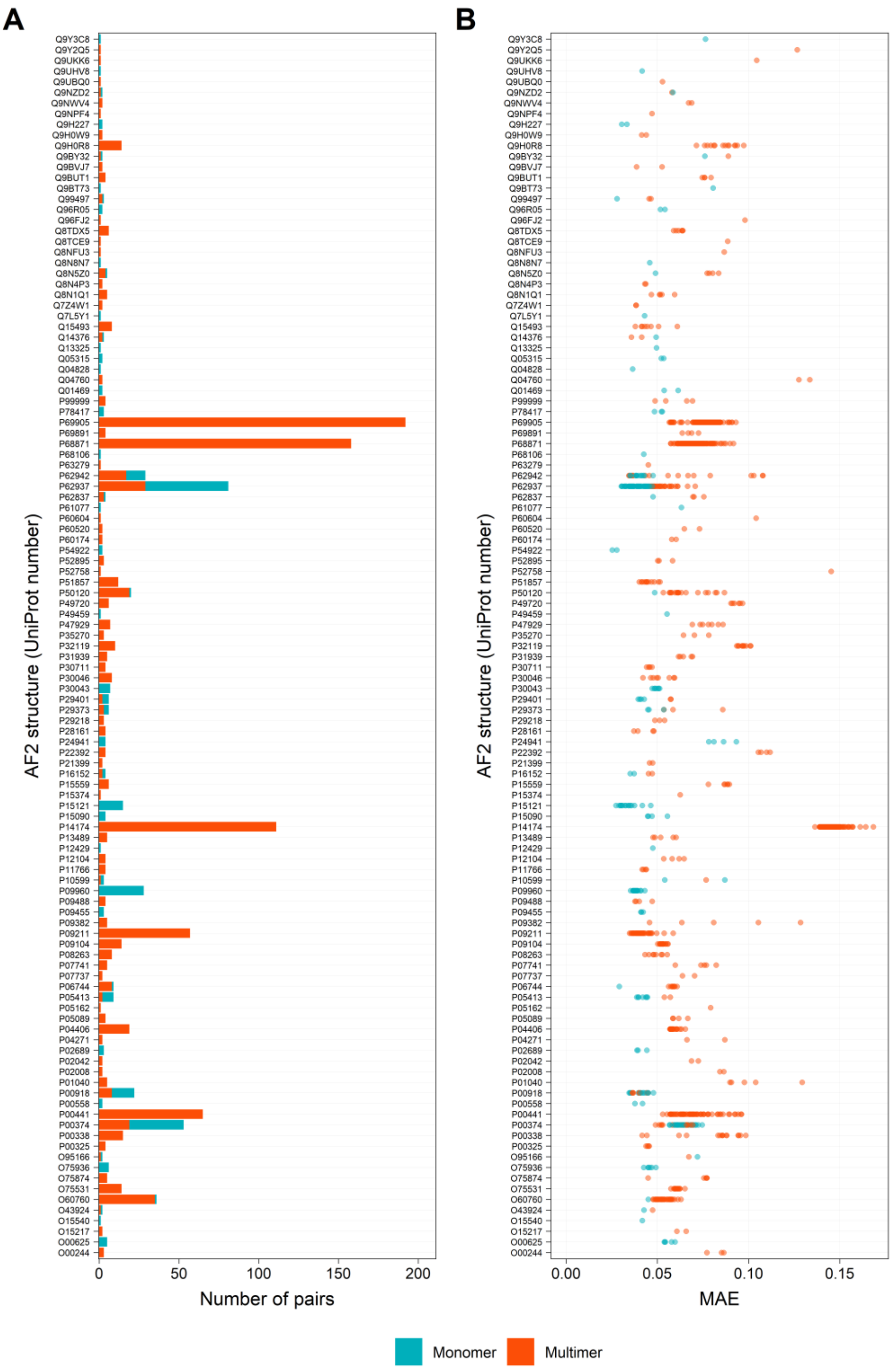


**Figure S3. Representation of each AlphaFold structure in the dataset. (A)** Number of matching experimental structures for each analyzed AlphaFold structure. The color indicates if the experimental structure is a monomer or a multimer. **(B)** Mean absolute error (MAE) for each structure pair. Each data point represents the MAE for one data pair. The pairs are organized according to unique AlphaFold structure. The color indicates if the experimental structure is a monomer or a multimer.

**Table S2. Error metrics as a function of pLDDT for each group of data pairs**

| Sequence overlap | Resolution | Mer | pLDDT | n | MAE | MSD | $SD_{Abs}$ |
|---|---|---|---|---|---|---|---|
| 100% | <=2.0 | Monomer | 20-30 | 1 | 0.522 | 0.522 | NA |
| 100% | <=2.0 | Monomer | 30-40 | 7 | 0.252 | 0.078 | 0.204 |
| 100% | <=2.0 | Monomer | 40-50 | 10 | 0.281 | 0.006 | 0.192 |
| 100% | <=2.0 | Monomer | 50-60 | 10 | 0.276 | 0.225 | 0.192 |
| 100% | <=2.0 | Monomer | 60-70 | 28 | 0.169 | 0.063 | 0.162 |
| 100% | <=2.0 | Monomer | 70-80 | 34 | 0.157 | 0.018 | 0.142 |
| 100% | <=2.0 | Monomer | 80-90 | 150 | 0.124 | 0.004 | 0.159 |
| 100% | <=2.0 | Monomer | 90-100 | 2927 | 0.036 | 0.001 | 0.062 |
| 100% | <=2.0 | Multimer | 40-50 | 1 | 0.025 | -0.025 | NA |
| 100% | <=2.0 | Multimer | 50-60 | 4 | 0.359 | 0.344 | 0.306 |
| 100% | <=2.0 | Multimer | 60-70 | 9 | 0.278 | 0.195 | 0.197 |
| 100% | <=2.0 | Multimer | 70-80 | 20 | 0.191 | 0.101 | 0.186 |
| 100% | <=2.0 | Multimer | 80-90 | 111 | 0.194 | 0.087 | 0.213 |
| 100% | <=2.0 | Multimer | 90-100 | 3411 | 0.063 | 0.029 | 0.119 |
| >99% <100% | <= 1.5 | Monomer | 40-50 | 1 | 0.054 | 0.054 | NA |
| >99% <100% | <= 1.5 | Monomer | 50-60 | 4 | 0.435 | 0.207 | 0.224 |
| >99% <100% | <= 1.5 | Monomer | 60-70 | 7 | 0.194 | -0.125 | 0.210 |
| >99% <100% | <= 1.5 | Monomer | 70-80 | 9 | 0.219 | 0.025 | 0.167 |
| >99% <100% | <= 1.5 | Monomer | 80-90 | 76 | 0.086 | -0.014 | 0.093 |
| >99% <100% | <= 1.5 | Monomer | 90-100 | 4116 | 0.041 | 0.001 | 0.068 |
| >99% <100% | <= 1.5 | Multimer | 40-50 | 1 | 0.743 | 0.743 | NA |
| >99% <100% | <= 1.5 | Multimer | 50-60 | 2 | 0.588 | 0.588 | 0.328 |
| >99% <100% | <= 1.5 | Multimer | 60-70 | 11 | 0.251 | 0.209 | 0.186 |
| >99% <100% | <= 1.5 | Multimer | 70-80 | 42 | 0.192 | 0.034 | 0.171 |
| >99% <100% | <= 1.5 | Multimer | 80-90 | 152 | 0.130 | 0.038 | 0.176 |
| >99% <100% | <= 1.5 | Multimer | 90-100 | 6041 | 0.051 | 0.020 | 0.100 |
| >99% <100% | >1.5 <=2.0 | Monomer | 50-60 | 3 | 0.218 | 0.083 | 0.208 |
| >99% <100% | >1.5 <=2.0 | Monomer | 60-70 | 15 | 0.182 | 0.031 | 0.123 |
| >99% <100% | >1.5 <=2.0 | Monomer | 70-80 | 34 | 0.155 | -0.008 | 0.162 |
| >99% <100% | >1.5 <=2.0 | Monomer | 80-90 | 232 | 0.129 | -0.024 | 0.147 |
| >99% <100% | >1.5 <=2.0 | Monomer | 90-100 | 7652 | 0.040 | 0.000 | 0.067 |
| >99% <100% | >1.5 <=2.0 | Multimer | 40-50 | 1 | 0.180 | 0.180 | NA |
| >99% <100% | >1.5 <=2.0 | Multimer | 50-60 | 12 | 0.265 | 0.119 | 0.178 |
| >99% <100% | >1.5 <=2.0 | Multimer | 60-70 | 17 | 0.244 | 0.137 | 0.219 |
| >99% <100% | >1.5 <=2.0 | Multimer | 70-80 | 50 | 0.239 | 0.138 | 0.283 |
| >99% <100% | >1.5 <=2.0 | Multimer | 80-90 | 382 | 0.131 | 0.047 | 0.161 |
| >99% <100% | >1.5 <=2.0 | Multimer | 90-100 | 14966 | 0.055 | 0.025 | 0.108 |

**Table S3. Error metrics as a function of $RSA_{Exp}$ for each group of data pairs**

| Sequence_overlap | Resolution | Mer | $RSA_{Exp}$ | n | MAE | MSD | $SD_{Abs}$ |
|---|---|---|---|---|---|---|---|
| 100% | <=2.0 | Monomer | 0-0.4 | 2560 | 0.035 | 0.010 | 0.072 |
| 100% | <=2.0 | Monomer | 0.4-0.8 | 585 | 0.098 | -0.005 | 0.103 |
| 100% | <=2.0 | Monomer | 0.8-1.2 | 153 | 0.130 | -0.072 | 0.185 |
| 100% | <=2.0 | Monomer | 1.2-1.6 | 9 | 0.392 | -0.275 | 0.385 |
| 100% | <=2.0 | Monomer | 1.6-2 | 2 | 0.034 | 0.034 | 0.027 |
| 100% | <=2.0 | Multimer | 0-0.4 | 3021 | 0.071 | 0.049 | 0.144 |
| 100% | <=2.0 | Multimer | 0.4-0.8 | 654 | 0.104 | 0.000 | 0.117 |
| 100% | <=2.0 | Multimer | 0.8-1.2 | 177 | 0.134 | -0.067 | 0.173 |
| 100% | <=2.0 | Multimer | 1.2-1.6 | 10 | 0.081 | 0.043 | 0.075 |
| >99% <100% | <= 1.5 | Monomer | 0-0.4 | 3370 | 0.032 | 0.005 | 0.059 |
| >99% <100% | <= 1.5 | Monomer | 0.4-0.8 | 776 | 0.096 | -0.007 | 0.101 |
| >99% <100% | <= 1.5 | Monomer | 0.8-1.2 | 203 | 0.111 | -0.043 | 0.144 |
| >99% <100% | <= 1.5 | Monomer | 1.2-1.6 | 10 | 0.090 | -0.001 | 0.065 |
| >99% <100% | <= 1.5 | Monomer | 1.6-2 | 1 | 0.126 | -0.126 | NA |
| >99% <100% | <= 1.5 | Multimer | 0-0.4 | 5256 | 0.054 | 0.032 | 0.116 |
| >99% <100% | <= 1.5 | Multimer | 0.4-0.8 | 1114 | 0.101 | 0.006 | 0.113 |
| >99% <100% | <= 1.5 | Multimer | 0.8-1.2 | 331 | 0.110 | -0.062 | 0.151 |
| >99% <100% | <= 1.5 | Multimer | 1.2-1.6 | 21 | 0.336 | -0.269 | 0.339 |
| >99% <100% | <= 1.5 | Multimer | 1.6-2 | 1 | 0.908 | -0.908 | NA |
| >99% <100% | <= 1.5 | Multimer | 2-2.4 | 1 | 1.027 | -1.027 | NA |
| >99% <100% | >1.5 <=2.0 | Monomer | 0-0.4 | 6466 | 0.033 | 0.007 | 0.058 |
| >99% <100% | >1.5 <=2.0 | Monomer | 0.4-0.8 | 1410 | 0.100 | -0.016 | 0.108 |
| >99% <100% | >1.5 <=2.0 | Monomer | 0.8-1.2 | 399 | 0.107 | -0.055 | 0.144 |
| >99% <100% | >1.5 <=2.0 | Monomer | 1.2-1.6 | 25 | 0.333 | -0.285 | 0.387 |
| >99% <100% | >1.5 <=2.0 | Monomer | 1.6-2 | 6 | 0.082 | -0.076 | 0.050 |
| >99% <100% | >1.5 <=2.0 | Multimer | 0-0.4 | 13258 | 0.059 | 0.040 | 0.128 |
| >99% <100% | >1.5 <=2.0 | Multimer | 0.4-0.8 | 2842 | 0.108 | 0.002 | 0.118 |
| >99% <100% | >1.5 <=2.0 | Multimer | 0.8-1.2 | 849 | 0.117 | -0.059 | 0.150 |
| >99% <100% | >1.5 <=2.0 | Multimer | 1.2-1.6 | 47 | 0.225 | -0.181 | 0.304 |
| >99% <100% | >1.5 <=2.0 | Multimer | 1.6-2 | 8 | 0.325 | -0.289 | 0.450 |

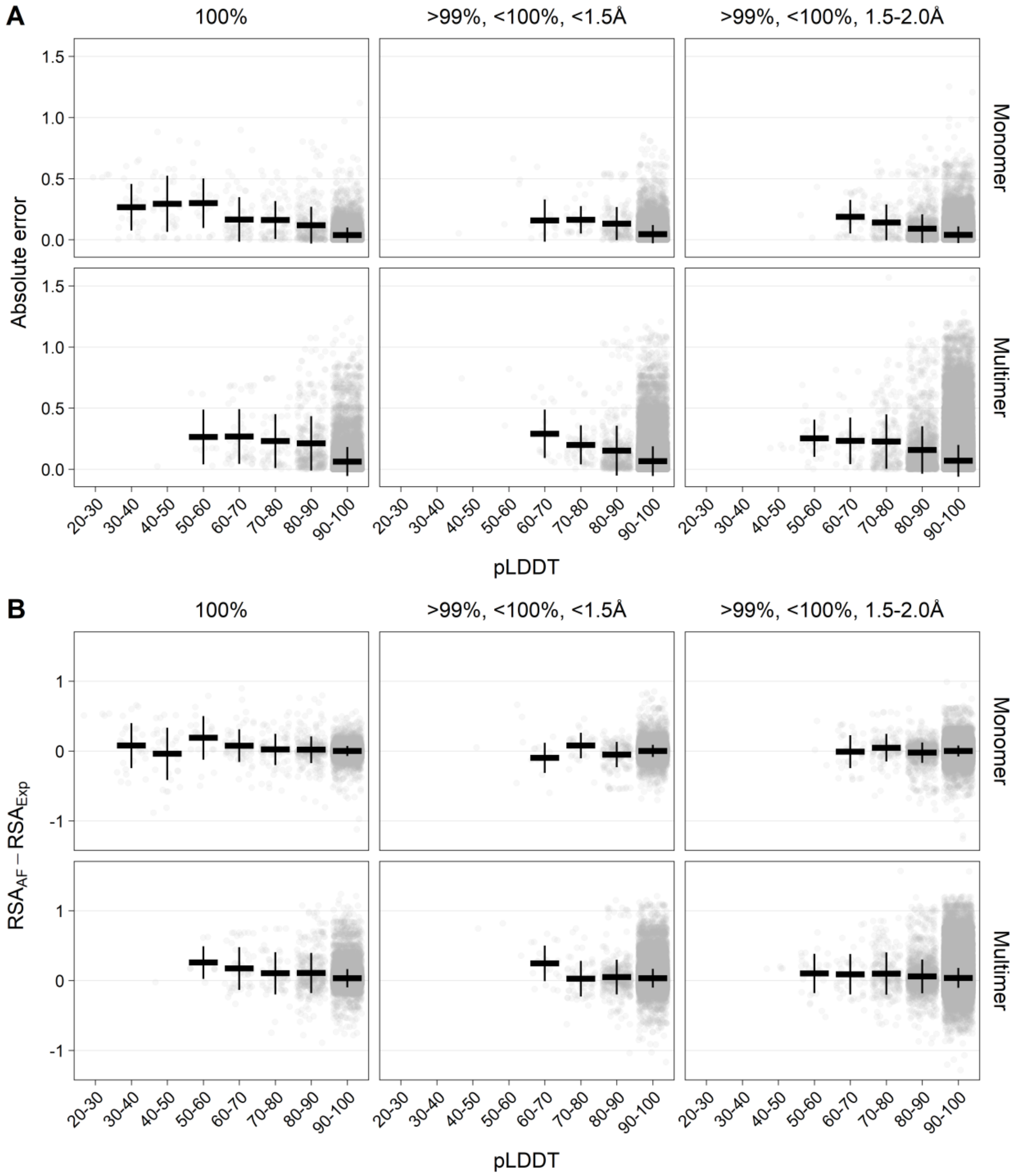


**Figure S4. Error vs pLDDT for six sub-datasets.** The pLDDT values are grouped in 10 percentage-point intervals, and gray points represent the actual data points, i.e., a point for each residue in a pair. Each panel shows residues from pairs that are grouped together based on sequence overlap, resolution of the experimental structure, and on the monomer-multimer status of the experimental structure. The six groups are disjointed from each other. **(A)** The MAE values (horizontal black bars) and one standard deviation ($SD_{Abs}$) above and below the MAE values (vertical black bars) are shown. MAE and $SD_{Abs}$ is not calculated for pLDDT groups with less than five data points. **(B)** The MSD values (horizontal black bars) and one standard deviation ($SD_{Signed}$) above and below the MSD values (vertical black bars) are shown. MSD and $SD_{Signed}$ is not calculated for pLDDT groups with less than five data points.

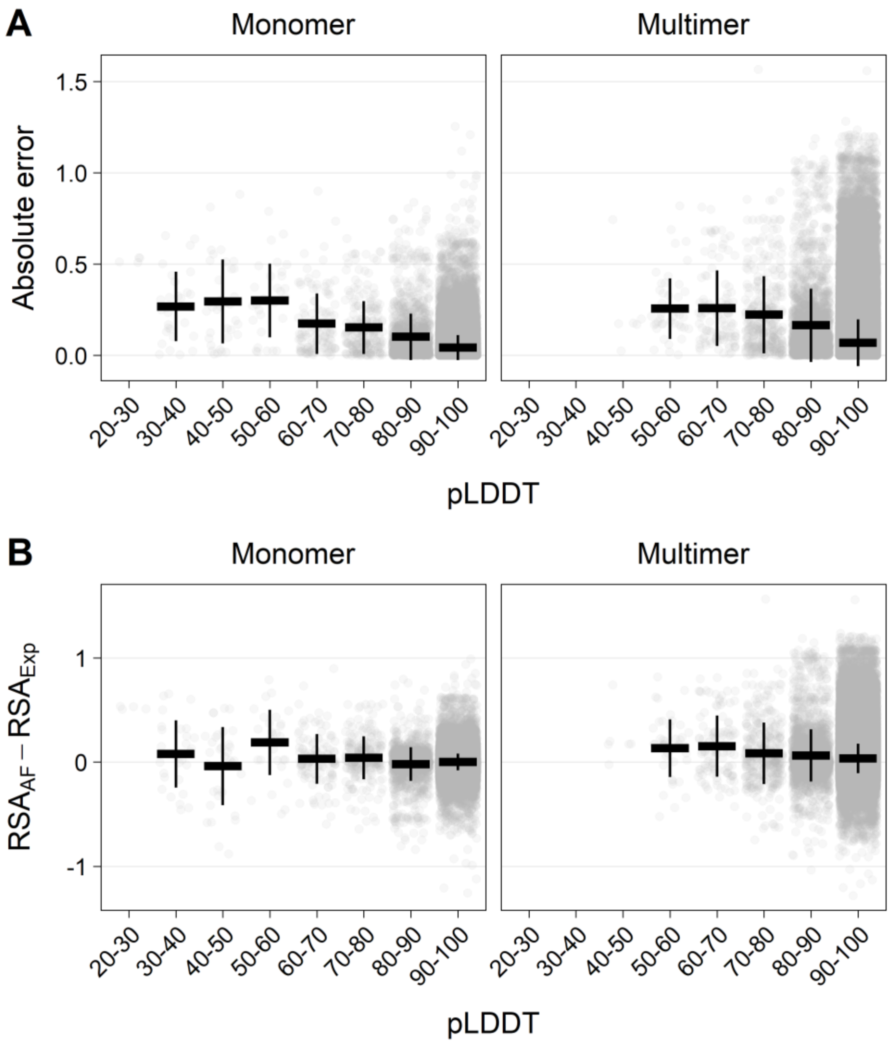


**Figure S5. Error vs pLDDT for monomer and multimer structures.** The pLDDT values are grouped in 10 percentage-point intervals, and gray points represent the actual data points, i.e a point for each residue in an experimental-AlphaFold pair. Each panel shows residues from pairs that are grouped together based on the monomer-multimer status of the experimental structure. **(A)** The MAE values (horizontal black bars) and one standard deviation ($SD_{Abs}$) above and below the MAE values (vertical black bars) are shown. MAE and $SD_{Abs}$ is not calculated for pLDDT groups with less than five data points. **(B)** The MSD values (horizontal black bars) and one standard deviation ($SD_{Signed}$) above and below the MSD values (vertical black bars) are shown. MSD and $SD_{Signed}$ is not calculated for pLDDT groups with less than five data points.

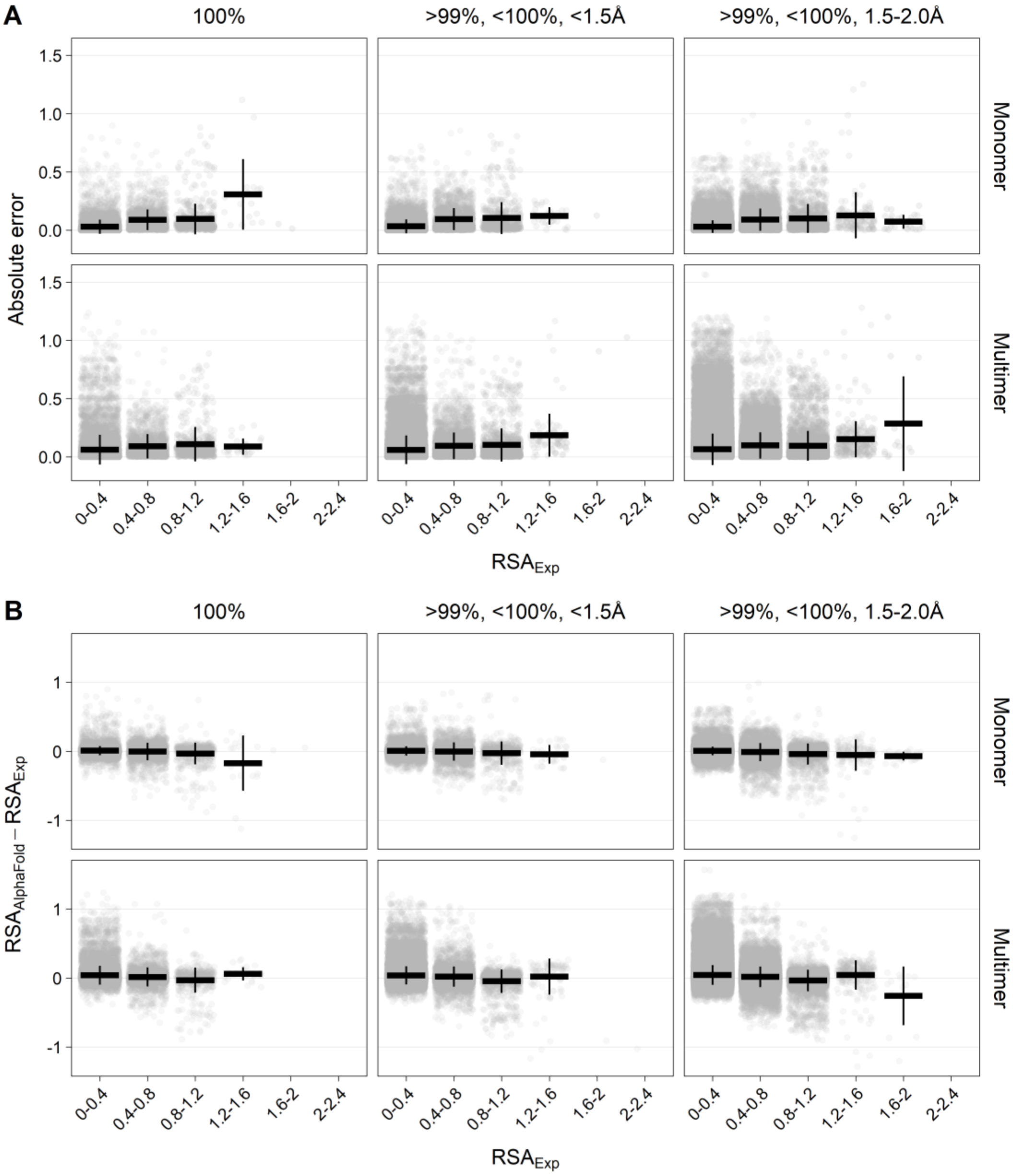


**Figure S6. Error vs experimental RSA for six sub-datasets.** The experimental RSA values are grouped in 0.4-intervals, and gray points represent the actual data points, i.e a point for each residue in an experimental-AlphaFold pair. Each panel shows residues from pairs that are grouped together based on sequence overlap, resolution of the experimental structure, and on the monomer-multimer status of the experimental structure. The six groups are disjointed from each other. **(A)** The MAE values (horizontal black bars) and one standard deviation ($SD_{Abs}$) above and below the MAE values (vertical black bars) are shown. MAE and $SD_{Abs}$ is not calculated for RSA groups with less than five data points. **(B)** The MSD values (horizontal black bars) and one standard deviation ($SD_{Signed}$) above and below the MSD values (vertical black bars) are shown. MSD and $SD_{Signed}$ is not calculated for RSA groups with less than five data points.

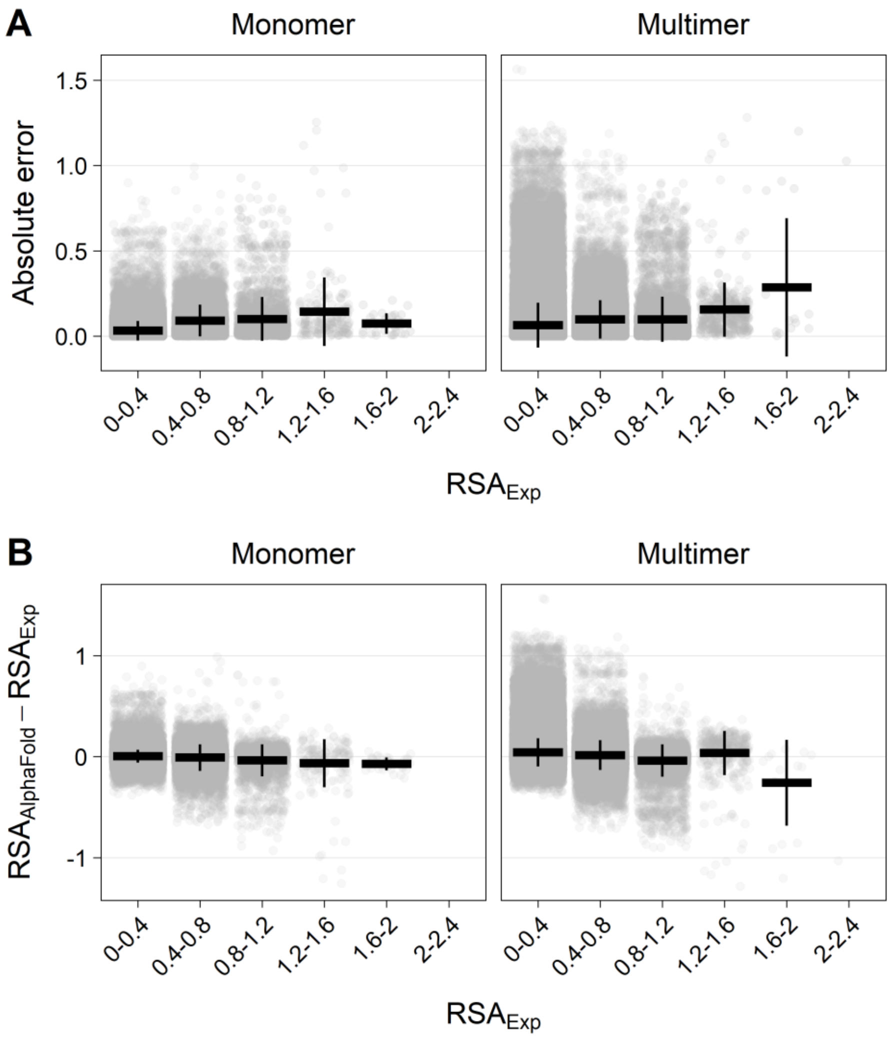


**Figure S7. Error vs experimental RSA for two monomers and multimers.** The experimental RSA values are grouped in 0.4-intervals, and gray points represent the actual data points, i.e a point for each residue in an experimental-AlphaFold pair. Each panel shows residues from pairs that are grouped together based on the monomer-multimer status of the experimental structure. **(A)** The MAE values (horizontal black bars) and one standard deviation ($SD_{Abs}$) above and below the MAE values (vertical black bars) are shown. MAE and $SD_{Abs}$ is not calculated for RSA groups with less than five data points. **(B)** The MSD values (horizontal black bars) and one standard deviation ($SD_{Signed}$) above and below the MSD values (vertical black bars) are shown. MSD and $SD_{Signed}$ is not calculated for RSA groups with less than five data points.

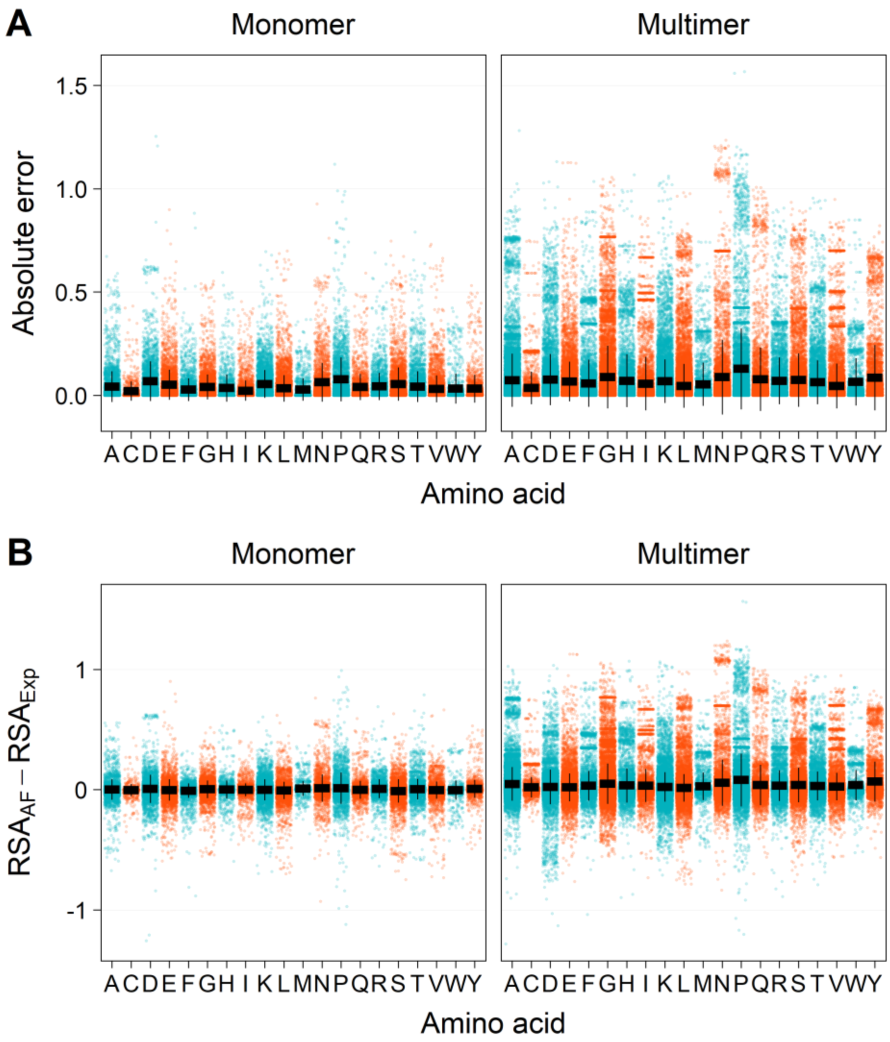


**Figure S8. Error vs amino acid type for two groups.** Red and blue points represent the actual data points, i.e a point for each residue in a data pair. Each panel shows residues from pairs that are grouped together based on the monomer-multimer status of the experimental structure. **(A)** The MAE values (horizontal black bars) and one standard deviation ($SD_{Abs}$) above and below the MAE values (vertical black bars) are shown. MAE and $SD_{Abs}$ is not calculated for amino acid types with less than five data points. **(B)** The MSD values (horizontal black bars) and one standard deviation ($SD_{Signed}$) above and below the MSD values (vertical black bars) are shown. MSD and $SD_{Signed}$ is not calculated for amino acid types with less than five data points.

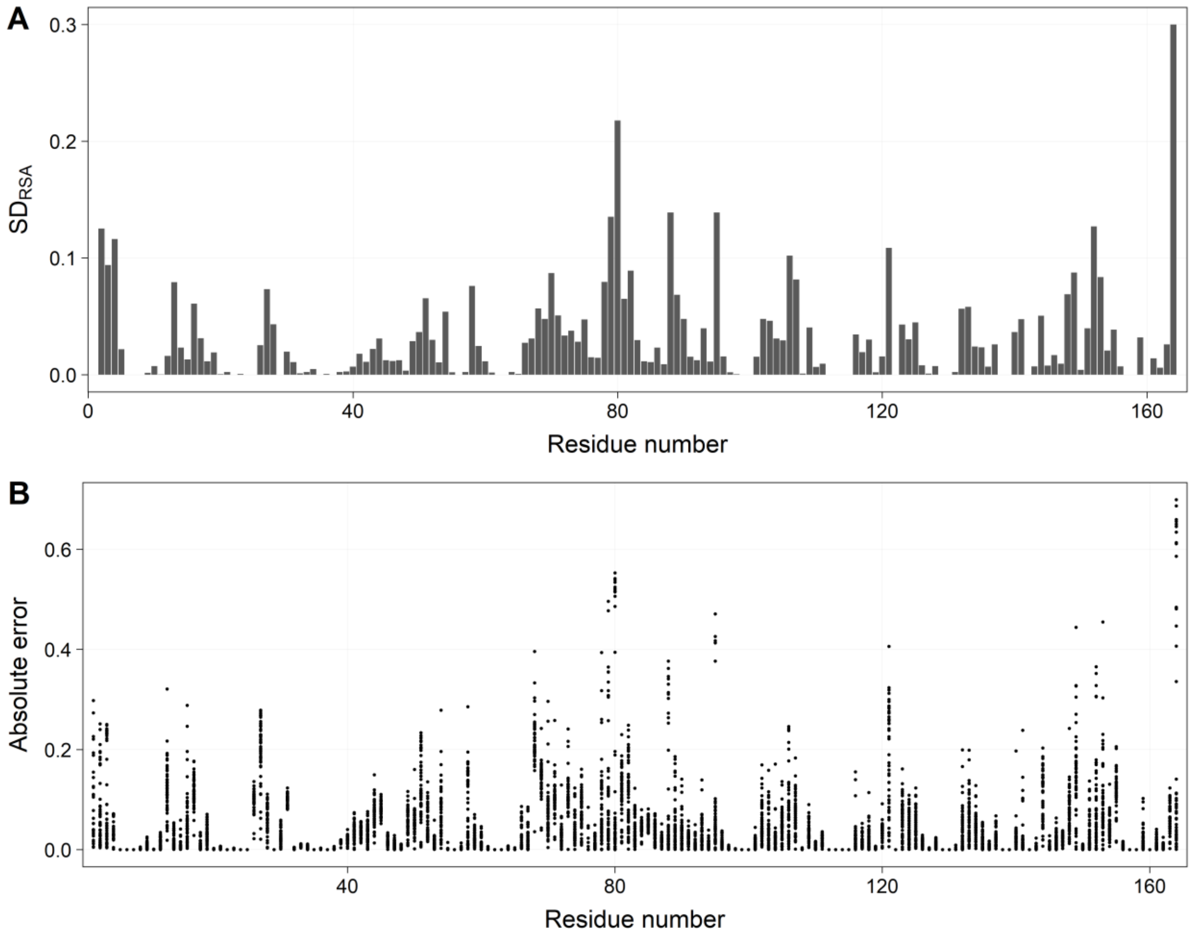


**Figure S9. Variation in $RSA_{Exp}$ and absolute error for peptidyl-prolyl *cis-trans* isomerase A (PPIA).** PPIA is matched to 52 monomeric experimental structures. **(A)** SD of $RSA_{Exp}$ for each residue in the chain. **(B)** Absolute error for each residue and each data pair.
